# 3D histology reveals that immune response to pancreatic precancers is heterogeneous and depends on global pancreas structure

**DOI:** 10.1101/2024.08.03.606493

**Authors:** Ashley L. Kiemen, Cristina Almagro-Pérez, Valentina Matos, Andre Forjaz, Alicia M. Braxton, Lucie Dequiedt, Jeeun Parksong, Courtney D. Cannon, Xuan Yuan, Sarah M. Shin, Jaanvi Mahesh Babu, Elizabeth D. Thompson, Toby C. Cornish, Won Jin Ho, Laura D. Wood, Pei-Hsun Wu, Arrate Muñoz Barrutia, Ralph H. Hruban, Denis Wirtz

**Affiliations:** Department of Pathology, Sol Goldman Pancreatic Cancer Research Center, Johns Hopkins School of Medicine, Baltimore, MD; Sidney Kimmel Comprehensive Cancer Center, Johns Hopkins School of Medicine, Baltimore, MD; Department of Chemical and Biomolecular Engineering, Johns Hopkins University, Baltimore, MD; Institute for NanoBioTechnology, Johns Hopkins University; Department of Functional Anatomy & Evolution, Johns Hopkins School of Medicine, Baltimore, MD; Bioengineering and Aerospace Engineering Department, Universidad Carlos III de Madrid, Leganés, Spain; Department of Pathology and Data Science Institute, Medical College of Wisconsin, Milwaukee, WI; Bioengineering Division, Instituto de Investigación Sanitaria Gregorio Marañón, Madrid, Spain

**Author notes:** Co-corresponding authors: Denis Wirtz, Ph.D. Croft 130, Department of Chemical & Biomolecular Engineering 3400 N Charles Street, Baltimore, MD 21218; Ashley Kiemen, Ph.D. Pathology 142 Department of Pathology 600 S Wolfe St Baltimore, MD 21287.

**Keywords:** Pancreatic cancer, PanIN, inflammation, tumor heterogeneity, precancers

## Abstract

Pancreatic ductal adenocarcinoma (PDAC) is a highly lethal cancer for which few effective therapies exist. Immunotherapies specifically are ineffective in pancreatic cancer, in part due to its unique stromal and immune microenvironment. Pancreatic intraepithelial neoplasia, or PanIN, is the main precursor lesion to PDAC. Recently it was discovered that PanINs are remarkably abundant in the grossly normal pancreas, suggesting that the vast majority will never progress to cancer. Here, through construction of 48 samples of cm^3^-sized human pancreas tissue, we profiled the immune microenvironment of 1,476 PanINs in 3D and at single-cell resolution to better understand the early evolution of the pancreatic tumor microenvironment and to determine how inflammation may play a role in cancer progression.

We found that bulk pancreatic inflammation strongly correlates to PanIN cell fraction. We found that the immune response around PanINs is highly heterogeneous, with distinct immune hotspots and cold spots that appear and disappear in a span of tens of microns. Immune hotspots generally mark locations of higher grade of dysplasia or locations near acinar atrophy. The immune composition at these hotspots is dominated by naïve, cytotoxic, and regulatory T cells, cancer associated fibroblasts, and tumor associated macrophages, with little similarity to the immune composition around less-inflamed PanINs. By mapping FOXP3+ cells in 3D, we found that regulatory T cells are present at higher density in larger PanIN lesions compared to smaller PanINs, suggesting that the early initiation of PanINs may not exhibit an immunosuppressive response.

This analysis demonstrates that while PanINs are common in the pancreases of most individuals, inflammation may play a pivotal role, both at the bulk and the microscopic scale, in demarcating regions of significance in cancer progression.

## INTRODUCTION

Invasive pancreatic ductal adenocarcinoma, or PDAC, is the most common form of pancreatic cancer and is projected to be the second leading cause of cancer-related deaths by 2030.^1^ While novel immunotherapies have made great strides in improving outcomes in malignancies including non-small cell lung cancer, melanoma, and Hodgkin’s lymphoma,^2–5^ immunotherapies are generally ineffective in pancreatic cancer.^6–8^ Paradoxically, inflammation is believed to play a key role in PDAC development and invasion.^9–11^ Individuals suffering from chronic pancreatitis have a 13-fold increase in risk of developing PDAC.^10^ Pancreatic cancer cells are surrounded by a dense network of fibrotic tissue containing immunosuppressive cells such as regulatory T cells, tumor-associated macrophages, and cancer-associated fibroblasts.^12–16^ As these stromal populations are believed to evolve early during pancreatic tumorigenesis,^11,17^ better understanding of the immune landscape in pancreata containing pancreatic cancer precursor lesions may improve our ability to develop effective strategies for immune-mediated cancer interception.

Pancreatic intraepithelial neoplasia (PanIN) is a noninvasive precursor lesion to PDAC that develops extensively throughout the pancreas with age.^18–20^ While most of us will develop PanINs, very few of these lesions will progress to invasive disease: the yearly incidence of PDAC diagnosed in the United States is roughly twelve per 100,000 individuals.^21–23^ Efforts to understand the drivers of PanIN initiation and progression have shown that the size, incidence, and genetic variation of these lesions is high.^18,20^ Here, we add to these efforts through in-depth quantitative 3D mapping and analysis of the immune microenvironment in exceptionally large (cm^3^) samples of human pancreas using CODA.

CODA is a workflow for quantitative 3D mapping of tissues using serially sectioned, hematoxylin and eosin (H&E) stained and digitized microscope slides that has been used extensively to study the 3D microanatomy, transcriptomic signatures, and 3D genetic heterogeneity of PanINs at single-cell resolution.^11,18,24–29^ As with existing serial sectioning workflows^30–37^, CODA is compatible with integration of multiple histological stains for study of structures that are difficult to detect using H&E alone.^36–39^

Here, through extension of CODA to analyze immunohistochemically (IHC) stained serial histological images, we explore at cellular resolution the spatial landscape of immune infiltration in large 3D pancreas tissues containing PanINs. We confirm that pancreas inflammation is strongly correlated with PanIN density. Around the PanINs, we find striking heterogeneity in leukocyte, T cell, and regulatory T cell density, with more regulatory T cells around larger PanINs compared to smaller PanINs. Using imaging mass cytometry (IMC), we profile the immune composition around PanIN immune hotspots, revealing that a minority of PanINs possess a distinct, immunosuppressive microenvironment that often surrounds regions of higher grade of dysplasia and acinar atrophy. By quantifying the spatial heterogeneity of these hotspots, we find that the immune cell density around a PanIN lesion decorrelates in a span of tens of microns, suggesting that immune infiltration cannot be well characterized through assessment of a 2D tissue section. In sum, this work reveals the importance of 3D analysis in revealing biologically significant events in cancer progression.

## RESULTS

### Three-dimensional reconstruction of the pancreatic immune microenvironment at single-cell resolution

We assessed 48 samples of grossly normal human pancreas tissue surgically resected in response to various pancreatic abnormalities (Fig 1A, <u>see methods</u>). Samples were serially sectioned, stained with H&E every third slide, and digitized.^18^ To characterize immune infiltration, two samples were selected for IHC staining (Fig 1B). Every third slide was stained with CD45 to label leukocytes. In one sample, the remaining sections were dual stained with CD3 / FOXP3 to label T cells and regulatory T cells.

**Fig 1.**
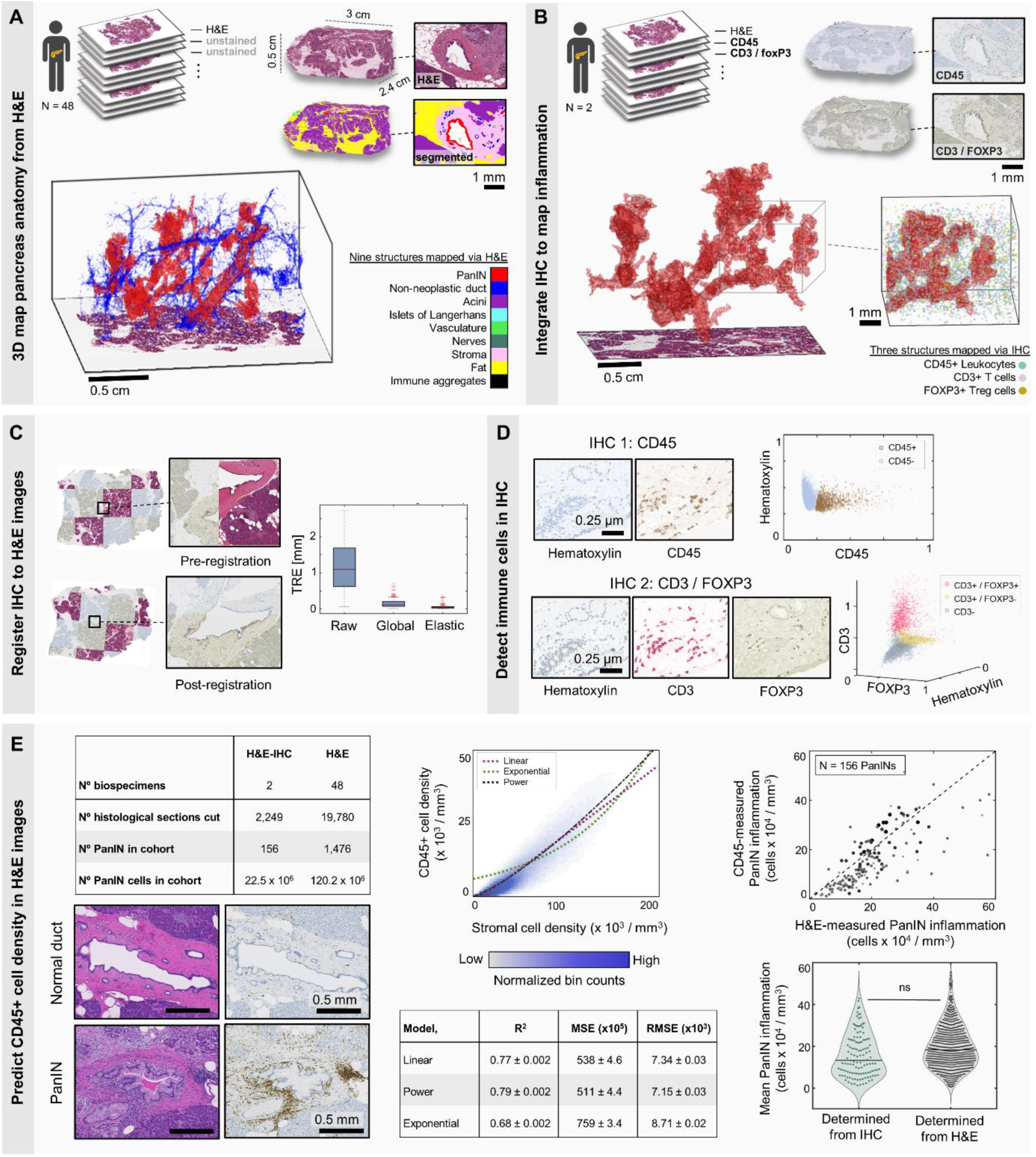
Integration of CODA + IHC for mapping the pancreatic immune microenvironment. (**a**) Forty-eight samples of cm^3^-sized human pancreas tissue were reconstructed using CODA to create 3D maps of pancreatic microanatomy. (**b**) In a subset of cases, intervening sections were stained for CD45 and CD3/FOXP3, enabling integration of immune cells in the 3D pancreas microenvironment. (**c**) Nonlinear image registration was used to align the multiplex images to the same coordinate space. Target registration error (TRE) demonstrates the quality of the registration. (**d**) Immune cell coordinates were generated using color deconvolution and a k-medoids algorithm. (**e**) Top left: table cataloging the number of tissue samples, sections, and PanIN per cohort. Bottom left: sample H&E and CD45-stained histology showing a non-inflamed normal pancreatic duct and an inflamed PanIN. Cell density in the stroma (pink in H&E image) appears to correlate to CD45+ stain. Center: approximation of 3D CD45+ cell density using 3D stromal cell density via five-fold cross-validation of linear, power, and exponential fits, with the best fit achieved using a power law. Top right: graph depicting error in approximation of mean PanIN inflammation using the power law fit. Bottom right: violin plot depicting the mean PanIN inflammation as determined using CD45-stained IHC images from two 3D samples and as determined using H&E images from 46 3D samples.

We adapted CODA nonlinear image registration for application to datasets with multiple stains (Fig 1C). CODA registers images by maximizing pixel cross-correlation and has been shown to outperform other registration algorithms in several parameters.^24^ In the case of registration of similarly stained image pairs (H&E to H&E or IHC to IHC), structures in the images are colored similarly (Fig S1A). However, for registration of multi-stained datasets, structures on adjacent sections may show variable staining patterns – for example collagen stains pink in H&E but is light grey in most IHC images, reducing pixel-to-pixel correlation. We tested three workflows to determine the optimal procedure: (1) registration of color images, (2) registration of the hematoxylin channel images, and (3) registration of the H&E images, followed by serial integration of the IHC images (Fig S1B). By comparing target registration error and warp, we found that the third method yielded the best results (Fig S1C).

We next used semantic segmentation^40^ to label ten structures in the pancreas at two µm resolution: PanIN, normal ducts, acinar tissue, islets of Langerhans, vasculature, nerves, fat, stroma, immune aggregates, and non-tissue (Fig 1A). When compared to an independent testing dataset, the model performed with an accuracy of 95.1% (FigS2A). Color deconvolution was used to de-mix the hematoxylin from the remaining stains and nuclear coordinates were generated.^24^ In IHC, the antibody signal in the area surrounding each nucleus was determined, and K-medoids clustering was used to distinguish CD45+ and CD45-cells, or CD3+ / FOXP3-, CD3+ / FOXP3+, and CD3- / FOXP3-cells (Fig 1D). Validation against manual counts revealed an average precision and recall of 0.89 and 0.88 (Fig S2B). Finally, as previously described,^24^ we adjusted our cell counts to account for over- or under-counting due to serial sectioning (Fig S2C). Integration of image registration, segmentation, and immune cell detection enabled 3D visualization of pancreas structures and the immune microenvironment.

After noting the high density of leukocytes in the pancreatic stroma, we hypothesized that we could accurately estimate CD45+ cell density from H&E-based measure of stromal cell density (Fig 1E). We extracted these data from the two samples containing H&E and IHC images. Using R^2^, mean squared error, and root mean squared error, we tested linear, power, and exponential fits to determine that a power law well approximated CD45+ cell density. To confirm that our model accurately estimated PanIN inflammation, we performed two checks. First, we compared IHC-measured to H&E-measured PanIN inflammation in the two samples containing H&E and IHC staining, revealing a close match. Second, we compared the IHC-measured PanIN inflammation in the two samples with H&E and IHC staining to the H&E-measured PanIN inflammation in the 46 samples with only H&E staining, revealing similar distributions between the two cohorts and demonstrating the accuracy of our power-law model.

Together, this workflow enabled us to reconstruct 48 cm^3^-sized samples of the pancreas immune microenvironment. Next, we assess correlations between pancreas structure and inflammation at microanatomical (µm) and bulk (cm) scale to map immune patterns around pancreatic precancers.

### PanINs are the most inflamed structures in most normal pancreases and possess complex immune patterns

We first broadly examined the 3D samples to identify and understand the major relationships between pancreatic structure and inflammation. We compared the bulk inflammation to patient demographics (Fig 2A), identifying significantly higher inflammation in samples surgically resected from the pancreatic head compared to those resected from the pancreatic tail. We found elevated but nonsignificant inflammation in the pancreases of individuals diagnosed with PDAC compared to those diagnosed with pancreatic neuroendocrine tumors (PanNETs), metastatic carcinomas from outside the pancreas, and other abnormalities necessitating pancreatic resection. We found no significant difference in inflammation between individuals based on age or sex.

**Figure 2.**
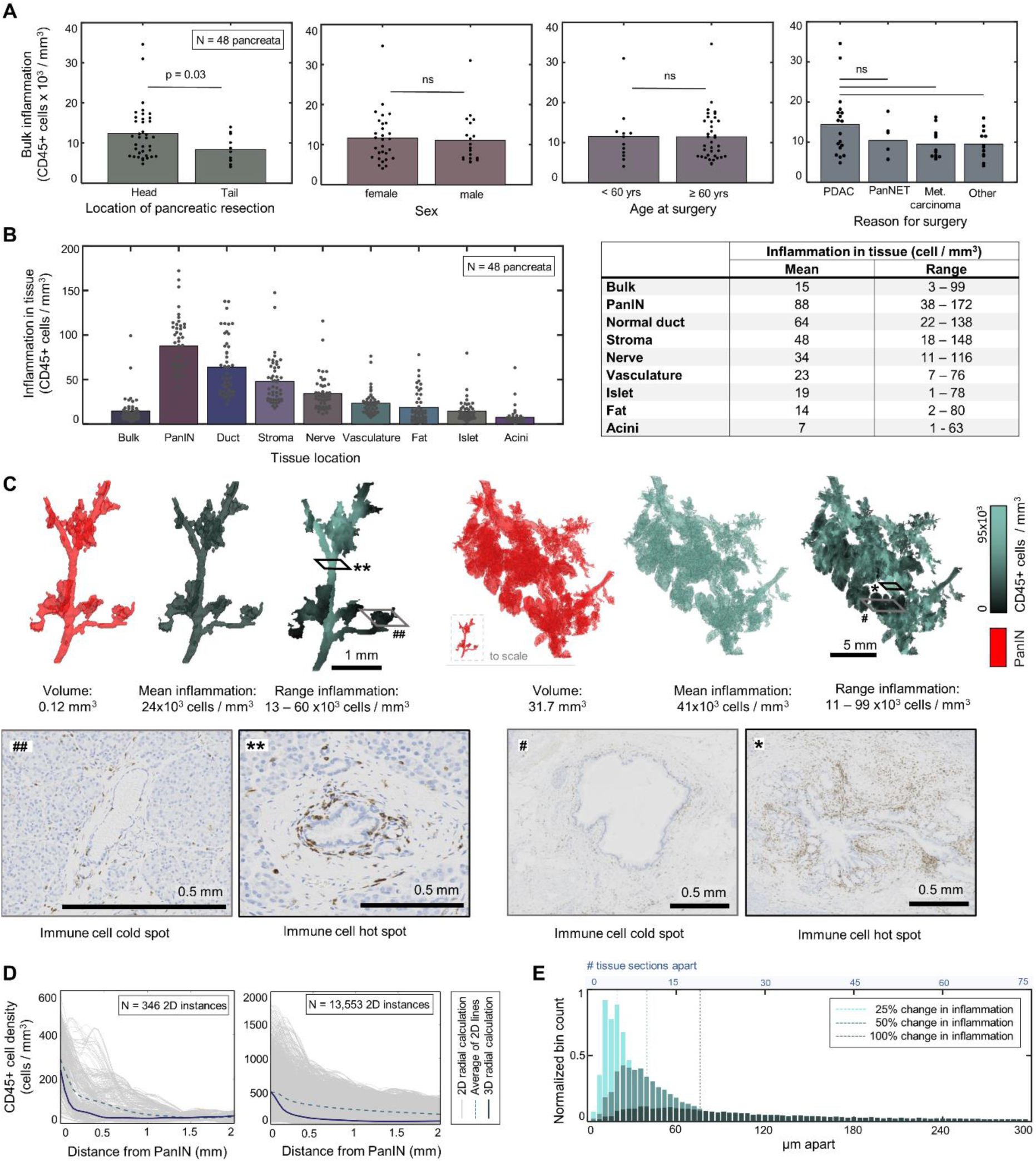
PanIN are most inflamed structure in most normal pancreases and feature complex immune patterns. (**a**) Samples resected from the pancreatic head were found to be significantly (p = 0.03) more inflamed than samples resected from the pancreatic tail. No significant difference in inflammation found as a function of patient diagnosis, age, or sex. P-values calculated using the Wilcoxon rank sum test. (**b**) Bar graph depicting the average inflammation present within eight structures of the pancreas. PanIN was found to be the most inflamed, followed by normal pancreatic ducts. Table containing mean and range values. (**c**) Sample 3D renderings depicting PanIN local immune cell density. Sample histology contains immune “hotspots” and “cold spots” present in these PanINs (**d**) Quantification of radial immune cell density around the PanINs depicted in **c**. Grey lines were calculated at each 2D instance of PanIN in the histology. Dashed blue line is the average of the 2D. Solid blue line is the true, 3D radial immune cell density. (**e**) Quantification of inflammatory heterogeneity. For a PanIN, 1,000 starting points were randomly chosen. Moving across the surface of the PanIN, the distance necessary for inflammation to change 25%, 50%, and 100% was found. On average, inflammation changed 25% within 5 sections (20 µm), 50% within 10 sections (40 µm), and 100% within 19 sections (76 µm).

We quantified the average immune cell density within eight structures of the pancreas to identify those most associated with inflammation (Fig 2B), defining local inflammation in a 150-µm radius around each structure. We found that PanIN lesions were often the most inflamed structure, followed by normal pancreatic ducts and stroma. Acinar lobules, islets of Langerhans, and fat were consistently the least inflamed structures, with the exception of one sample that contained extensive acinar to ductal metaplasia and so greater inflammation in the lobules (Fig S3A).

To understand the high immune cell density surrounding PanIN lesions, we generated 3D immunomaps (Fig 2C), where the 3D structure of the PanIN was overlaid by the local immune cell density. These immunomaps revealed that most PanIN lesions exhibit striking immune heterogeneity, possessing both immune “hotspots” and “cold spots.” We quantified this heterogeneity in immune cell density around PanINs in two ways: orthogonally away from the PanINs, and tangentially along the 3D external surface of the PanINs.

First, we compared the radial immune cell density around PanIN lesions in 2D and 3D (Fig 2D). In the graphs, each thin grey line represents the radial density around each 2D instance of the PanIN as it appears in the histology. The dashed blue line represents the bulk average of these 2D calculations, and the solid black line is the true, 3D radial immune cell density. This calculation demonstrates that random calculation of inflammation around a single 2D instance of a 3D structure can result in dramatic over- or under-estimation of the true immune response in a tissue, and that even the bulk average of 2D measurements fails to recapitulate true 3D information. Finally, we computed the “length scale” of inflammation at PanIN lesions, or the distance one would need to travel across the surface of a PanIN for the local inflammation to change by 25%, 50% or 100% (Fig 2E). We found that inflammation changes rapidly, on average changing 50% within 40µm, or 10 histological sections, suggesting that conclusions on the hot or cold nature of a tumor may be misleading if determined through assessment of a single 2D slide.

### Pancreatic inflammation correlates to pancreatic structure on bulk, but not microscopic, scale

We next investigated the hypothesis that higher incidence of PanIN correlates to higher levels of inflammation. We compared inflammation to pancreatic structure across microanatomical and bulk length scales (µm *vs*. cm scales). To compute the microanatomical correlation, we determined the cellular composition within the 150-µm radius around each of the 1,476 PanINs, as visualized in the sample 3D heatmap renderings (Fig 3A). To compute bulk correlation, we determined the 3D bulk cellular composition in each of the 48 3D pancreas samples, as visualized in the sample z-projections (Fig 3B). We determined the correlation coefficient of local inflammation to each local tissue structure, and similarly determined the correlation coefficient of bulk inflammation to each bulk tissue structure (Fig 3C).

**Figure 3.**
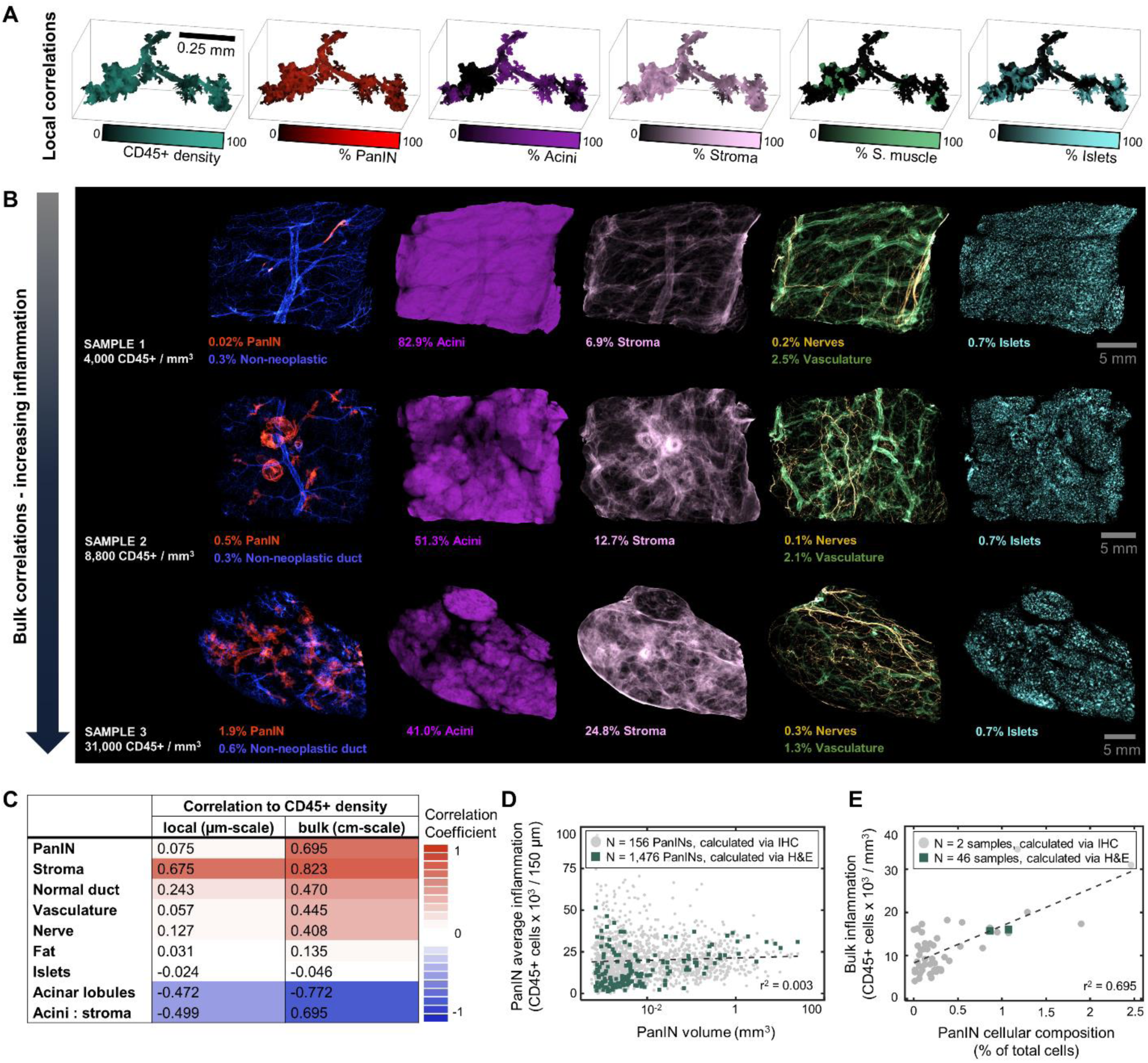
Pancreatic precancer inflammation is a global process. (**a**) 3D heatmap renderings of a PanIN depicting local composition of immune cells, and local density of pancreatic structures including stroma, acini, and vasculature. (**b**) Z-projections of three pancreatic tissue samples showing bulk changes to tissue structure with increasing PanIN content. (**c**) Table showing r^2^ values of the correlation between tissue structures and inflammation at the local (150 µm) scale and the bulk (entire cm^3^ tissue sample) scale. Several structures, including PanIN, stroma, normal ducts, vasculature, acinar lobules, and acinar-to-stromal ratio are highly correlated to inflammation at the bulk scale. (**d**) Average PanIN inflammation plotted as a function of PanIN volume. No correlation found. (**e**) Sample visualization of data in **c**, showing correlation of bulk inflammation to PanIN cellular composition.

Remarkably, while local inflammation correlated moderately with stromal and acinar composition, bulk inflammation correlated strongly with many pancreatic structures including stroma, acini, PanIN, normal duct, vasculature, and nerves. While locally we identify no correlation of PanIN volume to inflammation (Fig 3D), in bulk we identify strong correlation between overall PanIN cell fraction (PanIN cell number / total cell number in tissue) and inflammation (Fig 3E). These data suggests that while PanIN is highly correlated to inflammation, the associated immune cells are not always immediately surrounding the PanIN but may be elsewhere in the pancreatic parenchyma. This finding is supported by our identification of regions of highly inflamed acinar lobules that are upstream, but physically separated from large PanINs (Fig S3B).

### PanINs possess distinct immune hotspots and cold spots, characterized by differences in structure and immune cell content

We found that bulk pancreas inflammation is highly correlated to overall PanIN content, but that some of this inflammation surrounds the acinar lobules, not the PanIN themselves (Fig 3C). To better understand those immune cells that directly surround the PanINs, we next developed an algorithm to identify PanIN immune hotspots and cold spots. In each of the 48 3D samples, we pinpointed the coordinates of the ten hottest and ten coldest locations on a PanIN (Fig 4A). For each hotspot and cold spot region of interest (ROI), the algorithm output the corresponding H&E image from our serial histological dataset. Analysis of the histology revealed more high grade PanIN (PanIN-2 or PanIN-3), acinar to ductal metaplasia, and reactive stroma surrounding the PanIN hotspots, and pancreatic ducts containing large lumens or dilated pancreatic ducts associated with the PanIN cold spots. To confirm these results, we quantified the tissue composition in these regions of interest, revealing significantly higher composition of acini, islets of Langerhans, and PanIN in the histology outputs containing immune hotspots, and significantly higher composition of stroma and ductal lumen in the histology outputs containing immune cold spots (Fig 4B).

**Figure 4.**
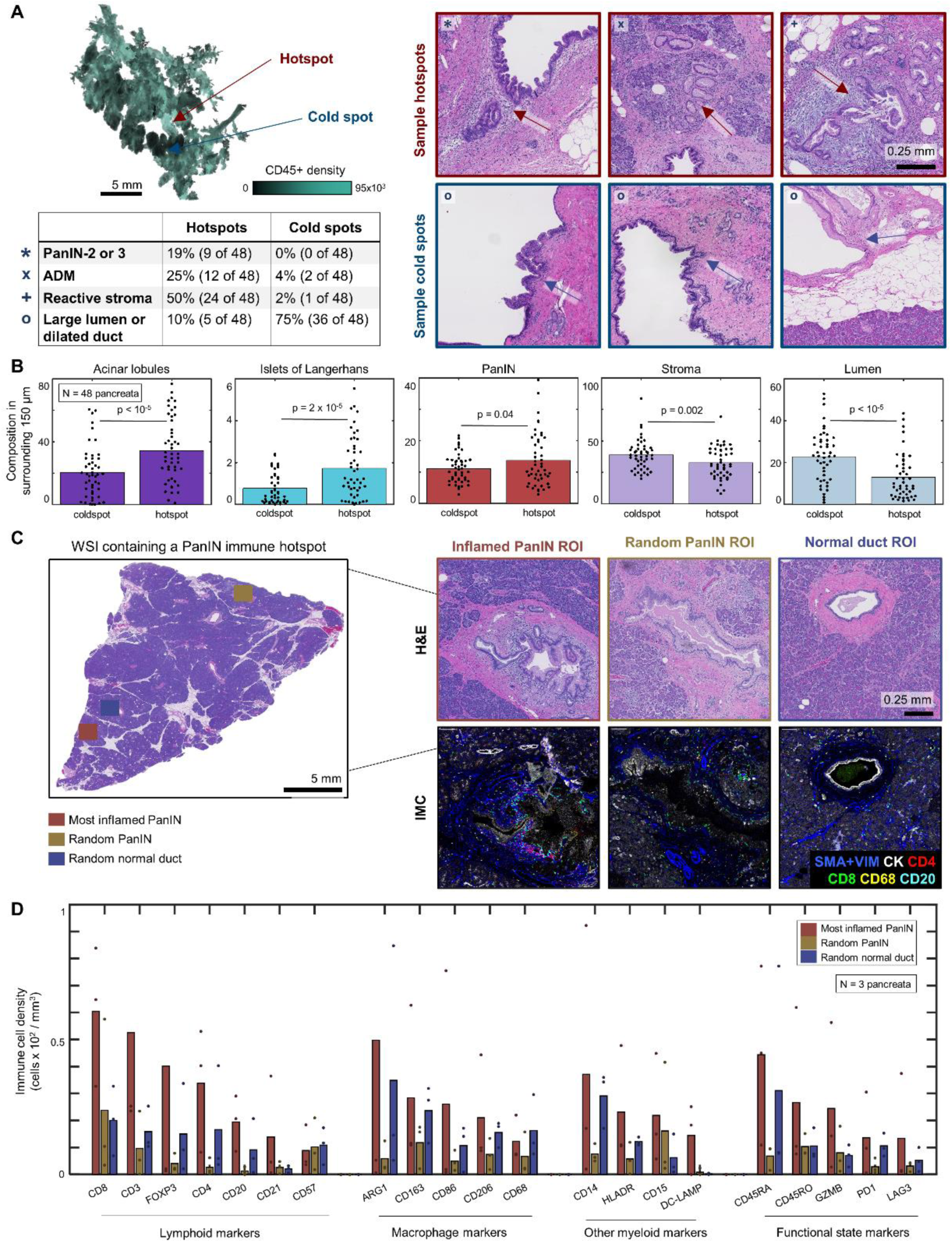
PanIN immune hotspots feature unique microenvironments. (**a**) A 3D rendering of a PanIN with regions of immune hotspots and cold spots. The table contains observed phenomena in the PanIN hotspots and cold spots for all 48 samples, with hotspots containing more high-grade dysplasia, reactive stroma, and acinar to ductal metaplasia (ADM), and cold spots containing more large ductal lumens or dilations. Sample histology provided, arrows indicate the regions of interest. (**b**) Comparison of the tissue composition in hotspot and cold spot histology revealed more acini, islets, and PanIN in hotspot regions, and more stroma and lumen in cold spot regions. P-values calculated using the Wilcoxon rank sum test. (**c**) In 3 samples, we identified a WSI containing a PanIN hotspot, another PanIN, and a normal pancreatic duct and applied a 38-plex imaging mass cytometry panel. (**d**) Quantitative comparison revealed generally higher immune cell densities at the hotspot PanIN. The immune cell densities of the randomly selected PanIN appeared to generally mirror those cell densities of the non-neoplastic duct.

To understand differences between PanIN immune hotspots and non-hotspot PanINs we applied an imaging mass cytometry (IMC) panel to label distinct immune cell types in a subset of three 3D pancreas samples (Table S1). We identified whole slide images in each sample containing a PanIN immune hotspot, a secondary, non-hotspot PanIN, and a non-neoplastic duct of similar radius and compared the IMC-derived immune cell densities around these regions (Fig 4C-D). We found generally higher density of all immune cell types around the PanIN hotspots. These hotspots were particularly notable for the relatively higher presence of pro-inflammatory cell types such as CD8+ cytotoxic T cells (along with markers of activation and/or exhaustion, e.g., Granzyme B, PD1, LAG3) and DC-LAMP+ dendritic cells, as well as immunosuppressive cell types such as foxP3+ regulatory T cells and macrophages (assessed by the expression of CD163, CD206, and Arg1). Remarkably, these data reveal that the immune profile around the non-inflamed PanIN lesions more closely resembles that of a non-neoplastic duct than that of the hotspot PanIN, suggesting that on average PanINs possess relatively low immune infiltrate, with the exception of distinct, hotspots that are easily missed without 3D analysis.

### PanIN size correlates to changes in presence of regulatory T cells

Finally, to understand changes to PanIN immune cell subtypes in 3-dimensions, we serially stained a single 3D pancreas sample alternately with H&E, CD45 to label leukocytes, and a dual stain to label CD3+ T cells and FOXP3+ regulatory T cells. This sample contained 34 spatially independent PanIN lesions. We created 3D immunomaps, revealing similar local cell densities between CD45+, CD3+, and FOXP3+ (Fig 5A). Confirming this observation, we computed the correlation coefficient between local CD45+, CD3+, and FOXP3+ cell density around all 34 PanINs in the 3D sample to find a high correlation between CD3 and CD45 and a moderate correlation between FOXP3 and CD3 (Fig S6A).

**Figure 5.**
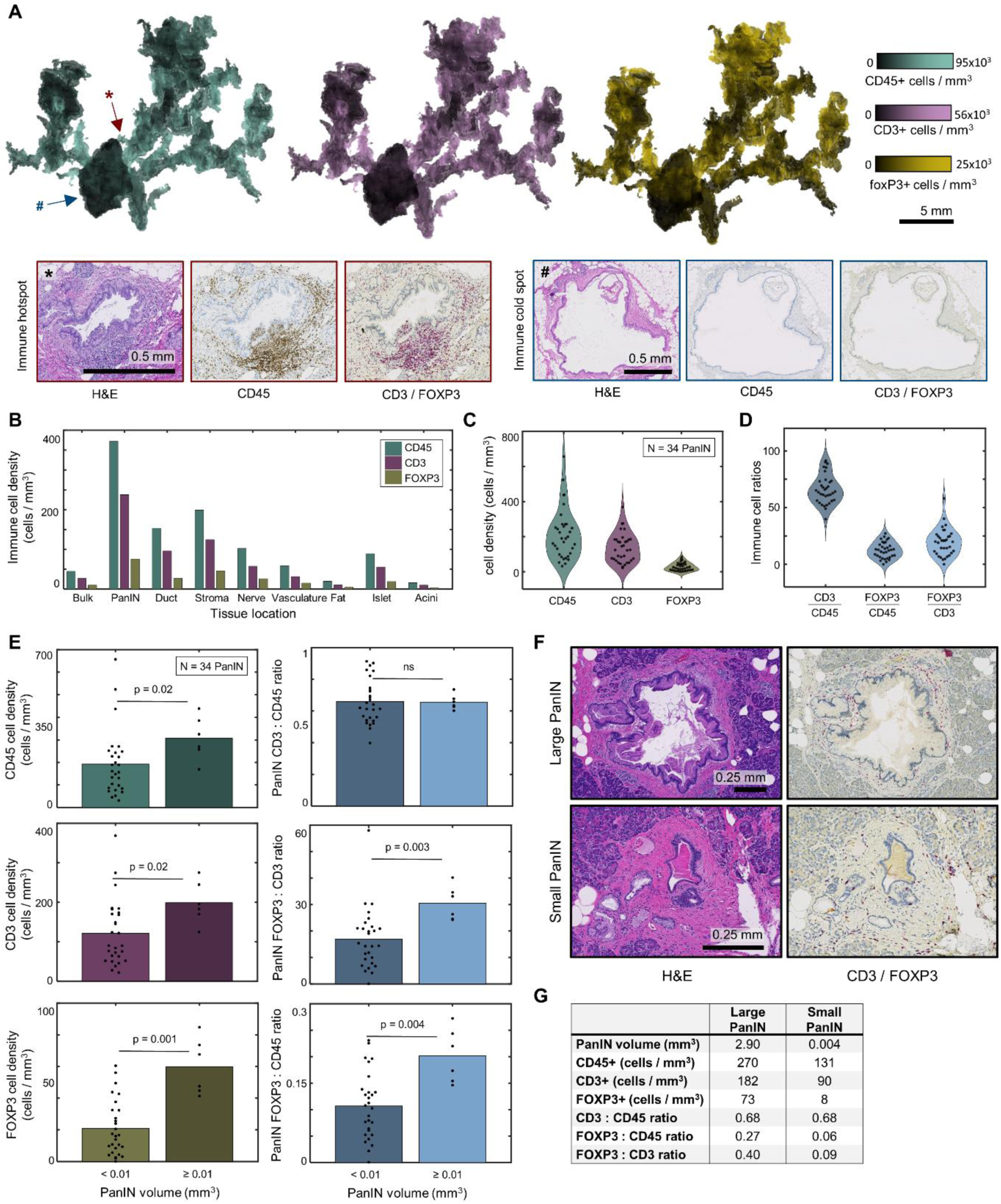
The composition of T cells around PanIN is heterogeneous. (**a**) 3D renderings of the CD45+, CD3+, and FOXP3+ cell density across a PanIN. Sample H&E and IHC histology at locations on the characterized by heavy or light immune infiltration. (**b**) Bar graph depicting the average CD45+, CD3+, and FOXP3+ cell density present within eight components of the pancreas. (**c**). Violin plot displaying the CD45+, CD3+, and FOXP3+ cell density at 34 PanIN. (**d**) Violin plot displaying the CD3 to CD45, FOXP3 to CD45, and FOXP3 to CD3 cell ratios at 34 PanIN. (**e**) Bar graphs depicting the CD45+, CD3+, and foP3+ cell density, and the CD3 to CD45, FOXP3 to CD45, and FOXP3 to CD3 cell ratios between small (< 0.01 mm^3^) and large (≥ 0.01 mm^3^) PanIN. Larger PanIN are in general more inflamed and have higher FOXP3 composition than small PanIN. P-values calculated using the Wilcoxon rank sum test. (**f**) Sample histology of a small PanIN possessing low FOXP3+ cell composition and a large PanIN possessing large FOXP3+ cell composition. (**g**) Table containing the immune cell properties of the PanIN shown in the sample histology.

We computed the density of immune cells in each structure, finding similar results to the calculation shown in Fig 2B, that PanIN is the most inflamed structure in the pancreas, followed by normal pancreatic ducts and stroma (Fig 5B).

Considering each of the 34 PanIN lesions in the 3D sample separately, we computed the volume of each PanIN, as well as its average CD45+, CD3+, and FOXP3+ cell density. Graphed in violin format, we find that PanIN lesions possess a range of immune cell densities (Fig 5C). For each PanIN, we similarly calculated the ratio of CD3+ to CD45+ cells, FOXP3+ to CD45+ cells, and FOXP3+ to CD45+ cells (Fig 5D).

Using 0.01 mm^3^ as a cutoff, we compared the immune cell density and composition between small and large PanINs (Fig 5E). We found that larger PanINs contain slightly higher densities of CD45+ cells and CD3+ cells, and much higher levels of FOXP3+ cells. Additionally, while the ratio of CD3+ to CD45+ was not related to PanIN size, we found significantly higher ratios of FOXP3+ to CD45+ cells and FOXP3+ to CD3+ cells in larger PanINs compared to smaller PanINs (bar plots in Fig 5E, scatter plots in Fig S6B-C). This suggests that regulatory T cells exist in higher density and make up a larger percentage of the total immune cells around more developed PanINs than they do around small PanINs. We show sample histology of a small and large PanIN (Fig 5F) possessing dramatically different immune cell densities and ratios. The larger PanIN contains a high ratio of regulatory T cells to T cells (ratio: 0.40) compared to the smaller PanIN (ratio: 0.09). Volumes and immune cell density information presented in the table in Fig 5G.

## DISCUSSION

In this work, we demonstrate the power of quantitative 3D mapping to reveal novel patterns in inflammation in early pancreatic carcinogenesis that span multiple length scales – from bulk changes to organ structure on the multi-cm scale to rapid shifts in local immune cell density on the µm scale.

We identified a strong correlation between bulk pancreas inflammation and several pancreatic structures including large-scale structures such as acinar lobular and stroma, as well as microscopic structures such as PanIN. We found that some of this inflammation appears surrounding the Panin lesions and some inflammation is further away, often located near upstream acinar atrophy and fibrosis (see again Fig S3B). This data provides strong evidence derived from human tissue samples to support proposed links between inflammation and neoplastic initiation in the pancreas, specifically the existence of a positive feedback loop between precancer abundance and pancreatic inflammation.^41–45^ The noted correlation between microscopic structures such as inflammation and PanINs to macroscopic structures such as pancreatic acinar lobules and stroma may explain the successes of works to non-invasively detect features such as pancreatic fibrosis, pancreatitis, lobulocentric atrophy, and PanINs using techniques such as MRI, CT, and EUS,^45–52^ and suggests that further development of imaging-based screening for pancreatic inflammation and PanINs is warranted. Extension of this 3D analysis to detailed assessment of samples from individuals diagnosed with chronic pancreatitis remains an important area for future study.

Additionally, we found that PanINs feature a heterogeneous immune response in 3D that is not well characterized through assessment of individual 2D histological slides. We found that the immunologically hottest regions on a PanIN are associated with higher-grade of dysplasia and acinar atrophy, and that the immunologically coldest regions on a PanIN are associated with large, open ductal lumens or dilations. Using imaging mass cytometry, we profiled the immune cell composition of these hotspots in comparison to ‘average PanIN in the cohort and to normal pancreatic ducts. We identified distinct immunological hotspots at PanINs rich in CD8+ T cells, regulatory T cells, cancer associated fibroblasts, and tumor associated macrophages, supporting previous work in human tissues suggesting these regions contain early immunosuppressive activity.^11^ In comparison, recent work profiled PanIN lesions in organ donor pancreata to find CD4+ T cells, fibroblasts and myeloid cells, but not regulatory T cells, at PanINs.^20^ We noted similar populations surrounding the ‘average’ PanIN regions in this cohort, but noted distinct, immunosuppressive cell types around the rarer, hotspot regions.

In sum, we find that pancreatic inflammation correlates strongly with pancreatic micro- and macro-structures, supporting the hypothesis of a positive feedback loop between inflammation and PanINs and supporting the feasibility of detecting pancreatic inflammation and PanINs in the clinic using non-invasive imaging. Through 3D assessment, we find that inflammation around PanIN lesions is heterogeneous, and that distinct hotspots composed of unique immunosuppressive cell types may hint at regions of the precancer that are at higher risk of progression to invasive cancer. This work demonstrates the ability of 3D mapping to quantify anatomical heterogeneity and to uncover rare biologically significant regions in large tissues.

## Data Availability Statement

The 3D rendering software used in this paper is available at the following GitHub page: https://github.com/ashleylk/CODA. Due to their large file size (TB scale per 3D sample), raw tissue data will be available from the corresponding authors upon request.

## Author Contribution Statement

DW, RHH, PHW, and LW conceived the project. AB, TCC, and JMB managed tissue collection, sectioning, staining, and scanning. ALK, CAP, VM, and LD completed all 3D methodological and analytic tasks. SS, AF JP, CDC, XY, and WJH assisted with the IMC collection and analysis. AMB, PHW, and ET advised on technical aspects of the project. ALK created the first draft of the manuscript and figures, which all authors edited and approved.

## Acknowledgements

The authors acknowledge the following sources of support: NIH/NCI T32 CA153952; NIH/NCI U54 CA268083; NIH UG3CA275681; S10OD034407; NIH/NCI U54 CA274371; PID2023-152631OB-I00; Sol Goldman Pancreatic Cancer Research Center; Lustgarten Foundation; Lustgarten Foundation-AACR Career Development Award for Pancreatic Cancer Research, in Honor of Ruth Bader Ginsburg; Rolfe Pancreatic Cancer Foundation; Susan Wojcicki and Denis Troper; The Joseph C. Monastra Foundation for Pancreatic Cancer Research; Fight Cancer Stay Positive Foundation; The Carl and Carol Nale Fund for Pancreatic Cancer Research.

## MATERIALS & METHODS

### Specimen acquisition and sample processing

This study was approved by the Institutional Review Board of the Johns Hopkins Hospital. Samples of grossly normal human pancreas tissue were harvested from the normal-adjacent tissue to a mass of clinical interest following surgical pancreatectomy. Tissues containing significant fibrosis, atrophy, or cancer in the normal adjacent region were excluded. The 48 samples analyzed here came from patients undergoing surgery for treatment of pancreatic ductal adenocarcinoma, pancreatic neuroendocrine tumors, serous cystadenomas, distal common bile duct adenocarcinomas, metastatic carcinomas from other organs, mucinous cystic neoplasms, tubulovillous adenomas of the duodenum, ampullary tumors, and lymphoepithelial cysts. The majority of these tissues were first described in a paper investigating the abundance and genetic heterogeneity of PanIN in grossly normal human pancreases.^18^

Samples were formalin-fixed, paraffin-embedded, and serial sectioned at a thickness of 4-microns. Every third section was stained with hematoxylin and eosin (H&E) and digitized at 20x magnification. In some samples, the intervening unstained slides were discarded or used for other purposes. In a subset of samples, the intervening tissue was cut onto plus slides, where it was integrated for labelling of immune cells using immunohistochemical (IHC) staining. Here, two large (>cm^3^) pancreas samples were 3D reconstructed using serial labelling of leukocytes using CD45. One of the samples also contains labels for T cells using CD3 and regulatory T cells using FOXP3. The remaining 46 samples contain serial H&E staining for mapping of gross anatomical pancreas structures and cell counts.

### CODA 3D reconstruction of pancreas microanatomy from H&E images

Samples were 3D reconstructed using CODA^24^, resulting in visualizable and quantifiable maps of human pancreas. The CODA workflow consists of four steps: image registration, nuclear detection, tissue multi-labelling, and visualization. Openslide software was used to save reduced size copies of all tissue images, corresponding to 2µm/pixel using nearest neighbor interpolation.^53^ For a pair of images, the registration was calculated through maximization of 2D cross correlation of pixel intensity to all images and correct for tissue rotation, translation, folding, splitting, and stretching. The CODA cell detection algorithm was used to quantify the cellularity of components via detection of 2D intensity peaks in the hematoxylin channel of the H&E images. Deep learning semantic segmentation was used to create microanatomical labels from histological images. Using annotations on a subset of histological images, the trained algorithm labelled, to a resolution of 2 µm, ten structures in histological images of the pancreas: islets of Langerhans, normal ductal epithelium, vasculature, fat, acinar tissue, collagen, PanIN, nerves, immune cell aggregates, and non-tissue whitespace with per class precision and recall of >90%. The image registration, cell detection, and tissue segmentation are integrated to create 3D reconstructions of pancreas microanatomy at large scale (up to multi-cm^3^), while maintaining cellular resolution.

### Registration of H&E and IHC images

For samples containing a mix of H&E and IHC images, the CODA image registration workflow was adapted to enable accurate and smooth integration of multi-plex datasets. Similar to the original CODA registration,^24^ sections were registered with a two-step process with the central image of each case serving as reference. Images were downsampled to a resolution of 8 µm / pixel. In a global rigid alignment step, the cross-correlation of a pair of whole slide images was maximized through determination of rotation and translation values. Then, in a local elastic step, images were cropped into patches (250 x 250 pixels) and rigid registration was applied to each region. Finally, displacement fields were interpolated from the grid of local registration data to generate maps to correct for non-uniform deformation between the images.

To optimize our H&E to IHC registration workflow, we tested three approaches: (1) registration of all raw, color images using the original CODA workflow, (2) registration of the hematoxylin channel of all images using the original CODA workflow, and (3) registration first of all H&E images using the original CODA workflow followed by serial integrative registration of the raw, color IHC images to the registered H&E images. The accuracy of registrations was assessed by calculating target registration error (TRE), change in tissue area from unregistered to registered images (Δarea), and maximum per-image magnitude of nonlinear displacement (tissue warp). Using 100 manually annotated fiducial landmarks on 50 pairs of images, TRE was calculated by determining the Euclidean distance between pairs of fiducial points in unregistered and registered images. To screen for shrinking or expanding of tissue sections caused by the registration process, Δarea was measured by calculating the %change in tissue area from unregistered to registered images. Finally, to screen for erroneous stretching of tissue regions during the nonlinear step of the registration process, the tissue warp was determined by calculating the difference between the maximum and minimum translations in the nonlinear displacement matrix. While this number is expected to be greater than zero (as zero means the image was only rigidly transformed), very large values imply non-biological stretching of the tissue.

### Detection of immune cell coordinates from IHC images

The previously described CODA cell detection algorithm was adapted to detect positive cells in IHC-stained images.^24^ First, the hematoxylin and antibody stain channels were isolated using color deconvolution. For each image, the pixels containing tissue were isolated by finding regions of low green channel intensity and high red-green-blue channel standard deviation. One hundred k-medoids clusters were calculated to represent the optical densities the tissue-containing pixels. The most common, blue-favored clusters were averaged to define the hematoxylin channel, and, for the CD45 stained images, the most common, brown-favored clusters were averaged to define the CD45 channel. The third channel was defined as the complement of the average of the hematoxylin and CD45 channels. For the dual stained CD3/FOXP3 stained images, the most common, red-favored clusters were averaged to define the CD3 channel, and the most common, black-favored clusters were averaged to define the FOXP3 channel. These optical densities were used to deconvolve the IHC images into their respective stains. To minimize variations in staining hue and saturation among images, histogram equalization was employed using the deconvolved channels of the first image of each sample as the reference. Next, CODA cell detection was used to generate nuclear coordinates from the hematoxylin channel of the images. The intensity of the antibody channels was determined at each nuclear coordinate. Joint histograms of the normalized intensities of the detected nuclei in the different channels were built. First, a joint histogram of the hematoxylin channel and the CD45 channel of the first IHC image of each sample was built to separate CD45+ and CD45-cells. Similarly, a joint histogram with three channels (hematoxylin, CD3, and FOXP3) of the first dual-IHC stained image was generated to separate T cells and Treg cells from the rest of cell types. In both cases, K-medoids partitioning^54^ was used to establish the separation thresholds.

To assess immune cell detection accuracy, four images of each CD45 stained images were randomly selected from each sample. Adjacent images were selected for the sample containing CD3 / FOXP3 staining. In this way, the same tissue region was assessed for all the immune cell types. From these images, tiles of 1 mm x 1 mm were extracted, containing hundreds of immune cells each. Cells were manually annotated. A manually labelled cell was considered equivalent to an automatically identified cell if the coordinates were within 5 µm of each other. The 5 µm was selected as the radius as this was determined to be half of the average radius of cytoplasm in the images. Finally, immune cell detection accuracy was assessed by measuring the precision and recall within each validation tile.

### Construction of 3D matrices for calculation of tissue metrics

The integration of image registration, cell detection, and tissue multi-labelling allowed construction of digital maps of tissue and cellular components. This reconstruction resulted in 3D matrices corresponding to each sample reconstruction. A 3D matrix containing tissue labels, a matrix containing nuclear coordinates generated from the H&E images, and a 3D matrix containing nuclear coordinates generated from the IHC images were obtained for the samples. Matrices were created at a resolution of 12×12×12 micron^3^. These matrices were used to perform all calculations shown in this manuscript. Calculations were performed in MATLAB 2022b.

### Calculation of volumes, cell counts, and cell densities

The volumes of each PanIN were calculated by summing the number of voxels in the tissue type matrix labelled as PanIN and converting from units of voxels to micron^3^. The number of cells of different structures were determined by dot multiplying the cell coordinate matrix by the tissue type matrix and summing the result. Cell counts determined via dot multiplication were corrected using the factors described in the cell detection section above. The number of immune cells surrounding a PanIN was determined using the tissue and cellular data matrices. The MATLAB function bwdist was used to determine the area of 150 microns surrounding each PanIN, and the number of immune cells located within that area was determined via dot multiplication of the distance matrix with the immune cell matrix.

### Calculation of local immune cell density and creation of 3D heatmap renderings

Local immune cell density was determined by counting the number of immune cells within a given distance of a voxel in the cell coordinates matrices and normalizing by the number of voxels labelled as tissue (and not whitespace) in the tissue type matrix. To visualize local immune cell densities, PanIN were first 3D rendered using the MATLAB functions isosurface and patch. The local immune cell density map was overlaid on this patch using a determined colormap (with black corresponding to low density values and a brighter color corresponding to high density values).

### Estimation of 3D CD45+ cell density from 3D stromal cell density

In two 3D samples containing H&E and IHC-stained sections, we compared the coordinates of detected CD45+ cells to those of all nuclei located in the H&E images. Initially, we calculated the immune cell density within local spheres of 150-micron radius. Next, the number of stromal cells was determined by dot multiplying the H&E cell matrix with a stromal mask. This mask was derived from the tissue type matrix label generated through deep learning 3D segmentation of each sample. The stromal cell density was then measured using the same 150-micron radius spheres.

From the two 3D pancreas matrices containing H&E and CD45+ cell data, we extracted CD45+ and stromal cell densities for 500,000 voxels each, resulting in a combined dataset of 1,000,000 points. Using this combined dataset of 1,000,000 voxels, we employed five-fold cross validation to evaluate linear, exponential, and power fits. We assessed the performance of these models using R2, mean squared error (MSE), and root mean squared error metrics. The resulting functions enabled the estimation of 3D CD45+ cell density based on 3D stromal cell density.

### Quantification of PanIN inflammatory heterogeneity

For all 1,476 PanIN in the 48 3D pancreas samples, 1,000 starting points were randomly chosen. Moving across the surface of each PanIN in 12 µm intervals, we measured the change in immune cell density with distance to determine the distance necessary for the inflammation to change 25%, 50%, and 100%. This was repeated for all 1,476 PanIN and plotted as a histogram.

### Calculation of 2D and 3D radial immune cell density

The MATLAB function bwdist was used to calculate the 2D or 3D radial distance of pixels in the tissue type matrix to a defined PanIN. For iterative distances away from the PanIN, the immune cell matrix was dot multiplied to determine the number of immune cells occupying a radial shell around the PanIN in 2D or 3D space. This number was normalized by the area or volume of the shell to determine the immune cell density at that distance from the PanIN. This calculation was performed starting at the external edge of the PanIN to a distance of 2 mm into the tissue. This radial data was visualized as a line (for calculation in 3D) or a series of lines (for calculation over a collection of 2D images) to show decay of immune cell density from the PanIN into the surrounding pancreas.

### Calculation of 3D local tissue density

Tissue density was determined by calculation of the number of voxels of a certain tissue type contained in a local sphere of 150-micron radius. This number was normalized by the number of tissue pixels (excluding nontissue whitespace) contained in the sphere. Correlation coefficient and p-values were determined using MATLAB 2022b.

### Detection of immune hotspots and cold spots around PanINs

Using the tissue label matrix, the location of all PanIN in each 3D sample was determined. The immune cell density at each PanIN voxel was computed through dot multiplication with the power-law adjusted stromal cell density matrix. The voxel with the highest immune cell density (a “hotspot”) was identified, and a 0.5 x 0.5 mm^2^ histological image centered on that coordinate was output, along with the tissue composition at that region of interest. All voxels within 0.5 mm of the hottest location in 3D space were eliminated, such that the next hotspot would be a minimum of 0.5 mm away from any previous hotspot. The next hottest location was found, and this process was repeated until ten hotspots were output. The process was repeated to identify the ten PanIN locations with the lowest immune cell density (“cold spots”) in each 3D sample.

### IMC staining and acquisition

Slides were cut from the FFPE blocks onto slides. The slides were first baked for two hours at 60°C, dewaxed in xylene wash, and rehydrated in an alcohol gradient (100%, 95%, 80%, 70% EtOH in Maxpar® H2O). The slides were then washed with Maxpar® Water, then incubated in an Antigen Retrieval Agent (Dako) at a temperature of 105°C for 1 hour. Slides were blocked with 3% BSA in Maxpar® PBS for 45 minutes. Selected antibodies were conjugated in-house, diluted to a concentration ranging from 0.25 mg / mL to 0.5 mg / mL, then aliquoted for use. The slides were stained with the final antibody cocktail (detailed in Table S1) overnight at 4°C. Slides were subsequently washed with Triton-X in Maxpar® PBS, then washed in Maxpar® PBS. For DNA labelling, Cell-ID™ Intercalator-Ir (Fluidugm) was diluted at 1:400 in Maxpar® PBS and stained. As a tissue counterstain, Ruthenium tetroxide 0.5% Stabilized Aqueous Solution (Polysciences) was diluted at 1:2000 in Maxpar® PBS and stained. A final wash was performed in Maxpar® Water. Images were acquired with a Hyperion Imaging System (Standard BioTools) at the Johns Hopkins Mass Cytometry Facility. Following acquisition, stacks of multi-layered ome.tiff images were exported, and representative images were generated utilizing MCD Viewer™ (Standard BioTools).

### Immune cell mapping using imaging mass cytometry

To quantify the resulting images, each channel of the ome.tiff file was exported as a separate TIF file. First, nuclear coordinates were generated using the CODA cell detection algorithm on the DAPI channel. Next, the intensity of each antibody channel was measured at each nuclear coordinate. Nuclei with an intensity greater than 50% of the maximum intensity were counted as positive cells. To exclude non-periductal stroma region, image mask were manually generated for each region of interest. The immune cell counts were converted to immune cell density through normalization by the periductal stroma area in mm2. Immune cell densities were compared across conditions (inflamed PanIN ROIs, randomly selected PanIN ROIs, and normal duct ROIs) using a bar graph.

---

**Fig S1.**
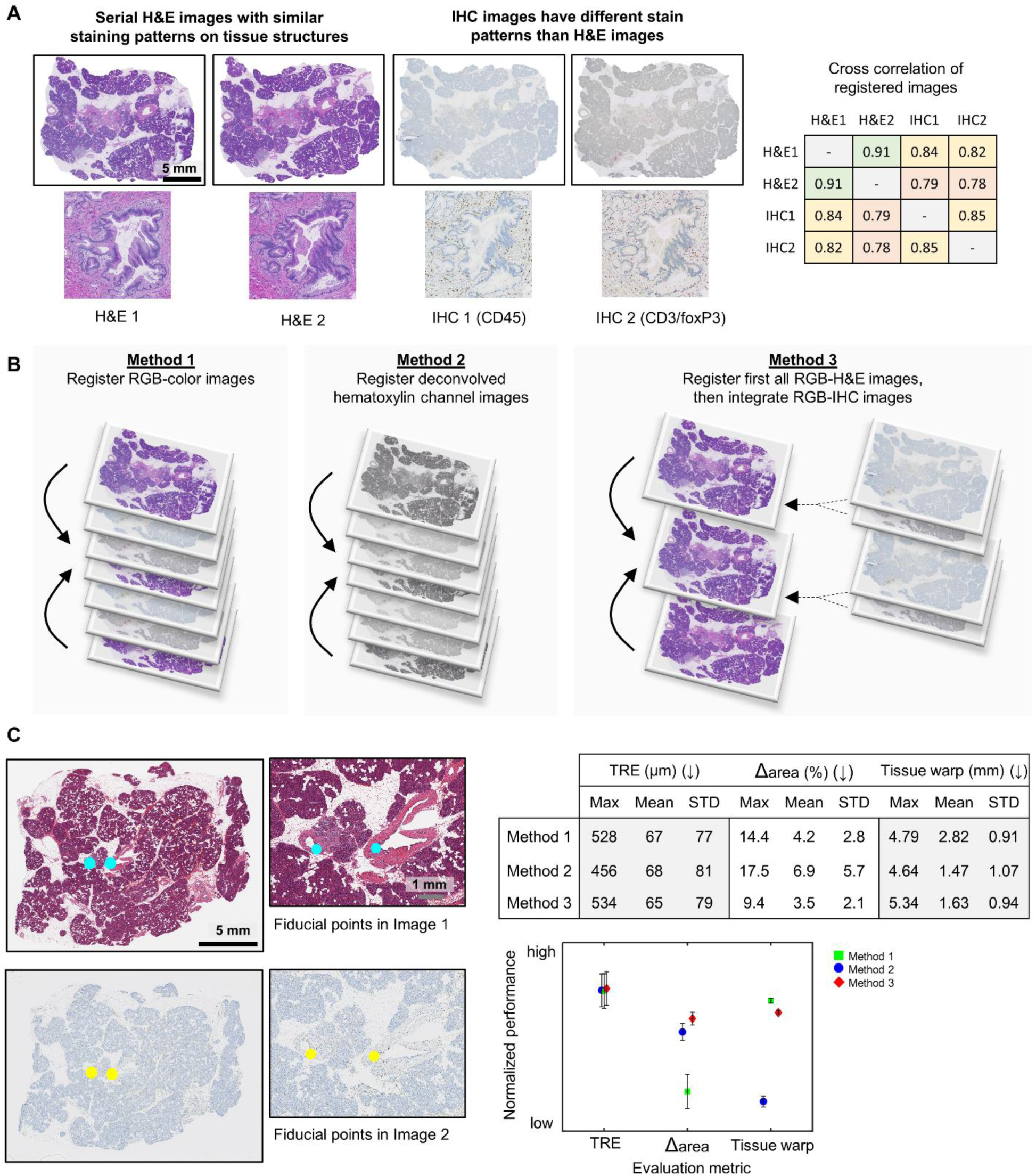
Validation of multiplex image registration workflow. (**a**) Left: sample H&E and IHC images display varying color intensities. Right: these differences are quantified using pixel-to-pixel 2D cross-correlation. (**b**) Three techniques for image registration of multiplex histological images: (left) co-register all color images, (center) co-register the extracted hematoxylin channel of all images, (right) first register H&E color images, then integrate IHC color images. (**c**) Left: manual generation of fiducial points between serial, multiplex images allowed quantification of each method’s performance. Top right: target registration error (TRE), change in tissue area (Δarea) and tissue warping were calculated for each method, showing an overall best performance of method 3. Bottom right: normalized, graphical representation of the registration validation metrics shown in the table.

**Fig S2.**
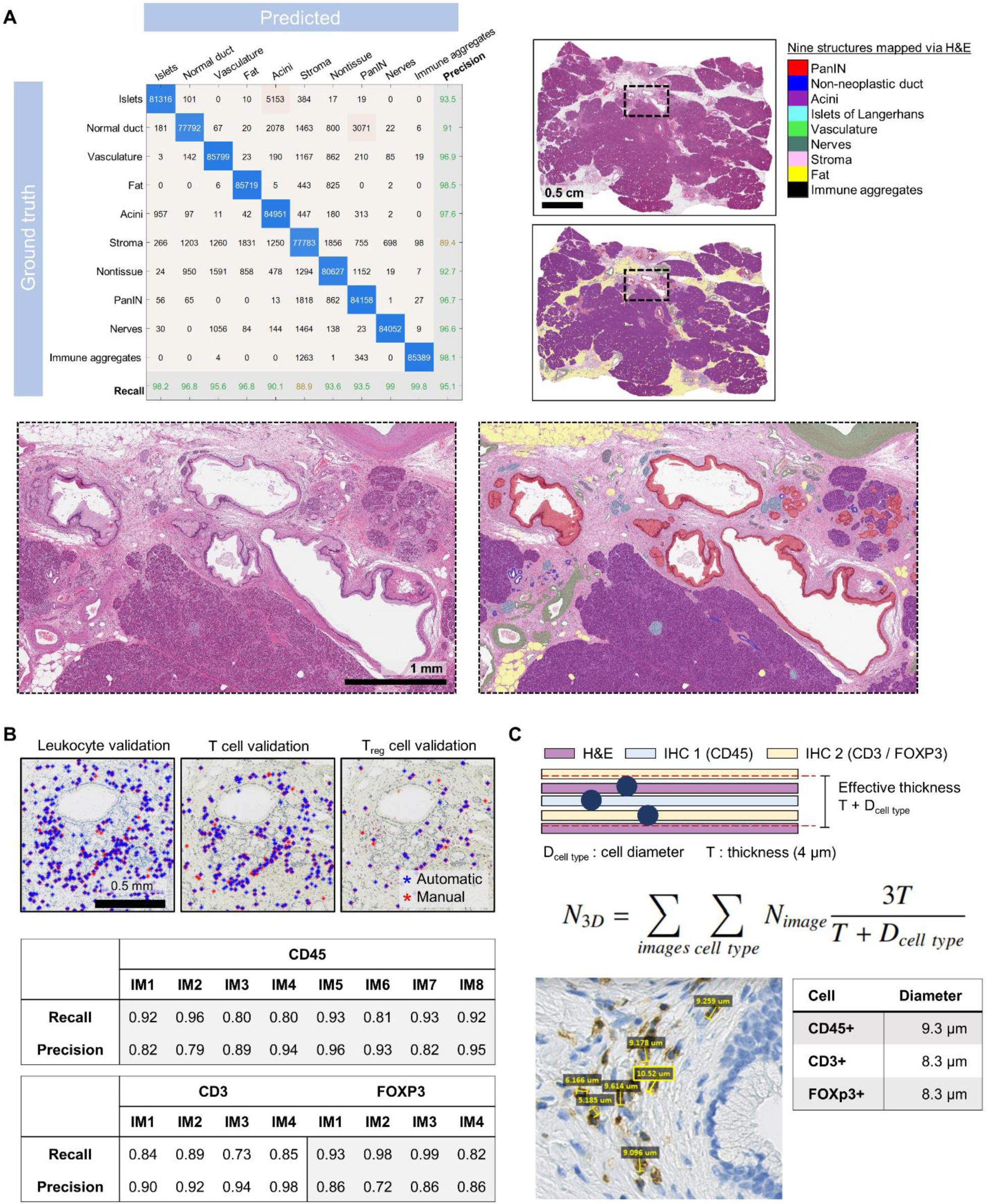
Validation of CODA tissue segmentation and immune cell detection from IHC. (**a**) Left: confusion matrix of semantic segmentation performance, with an overall accuracy of 96.6%. Right and bottom: sample human pancreas histology and segmented mask. (**b**) Top: sample histological sections used to validate detection of leukocytes, T cells, and regulatory T cells from IHC. Bottom: computed recall, precision, and F1 score. (**c**) The diameter of the IHC labelled immune cells was measured to extrapolate true 3D cell count from subsampled, serial histological images using the formula provided.

**Fig S3.**
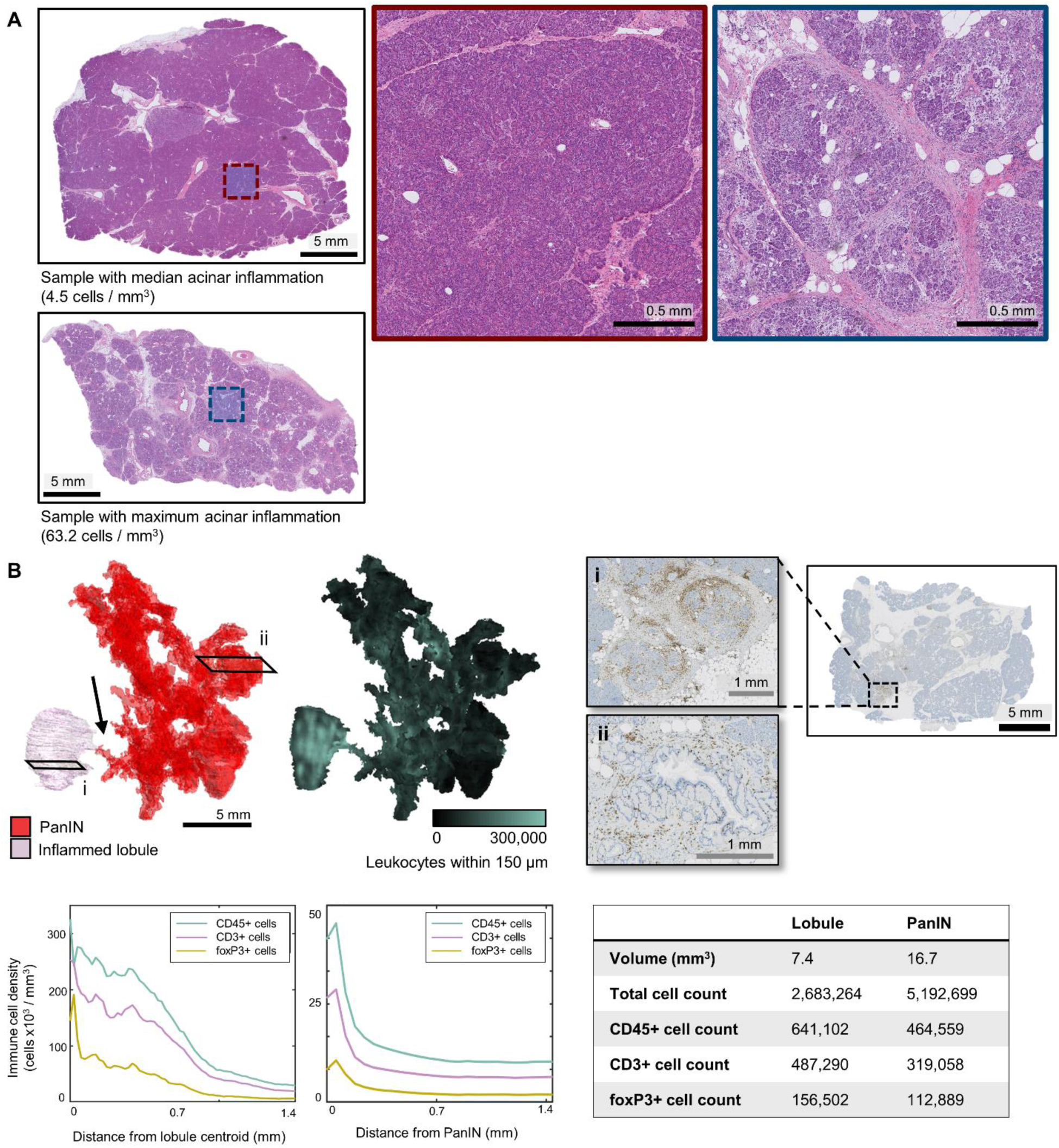
Immune aggregates associated with PanIN are sometimes distant. (**a**) Top and left crop: sample histology of a pancreas containing the median acinar immune cell density (4.5 CD45+ cells / mm^3^). Bottom and right crop: sample histology of a pancreas containing the maximum acinar immune cell density (63.2 CD45+ cells / mm^3^). The sample with higher inflammation in the acini contains extensive acinar to ductal metaplasia. (**b**) 3D renderings show a large PanIN located in a pancreatic duct directly upstream of a location of lobulocentric atrophy (black arrow indicates connection to duct). This region is highly inflamed, showing 6-fold greater local inflammation than the precursor lesion itself. Radial immune profiles around the PanIN and inflamed lobule are plotted, and bulk inflammation is shown in tabular form.

**Fig S4.**
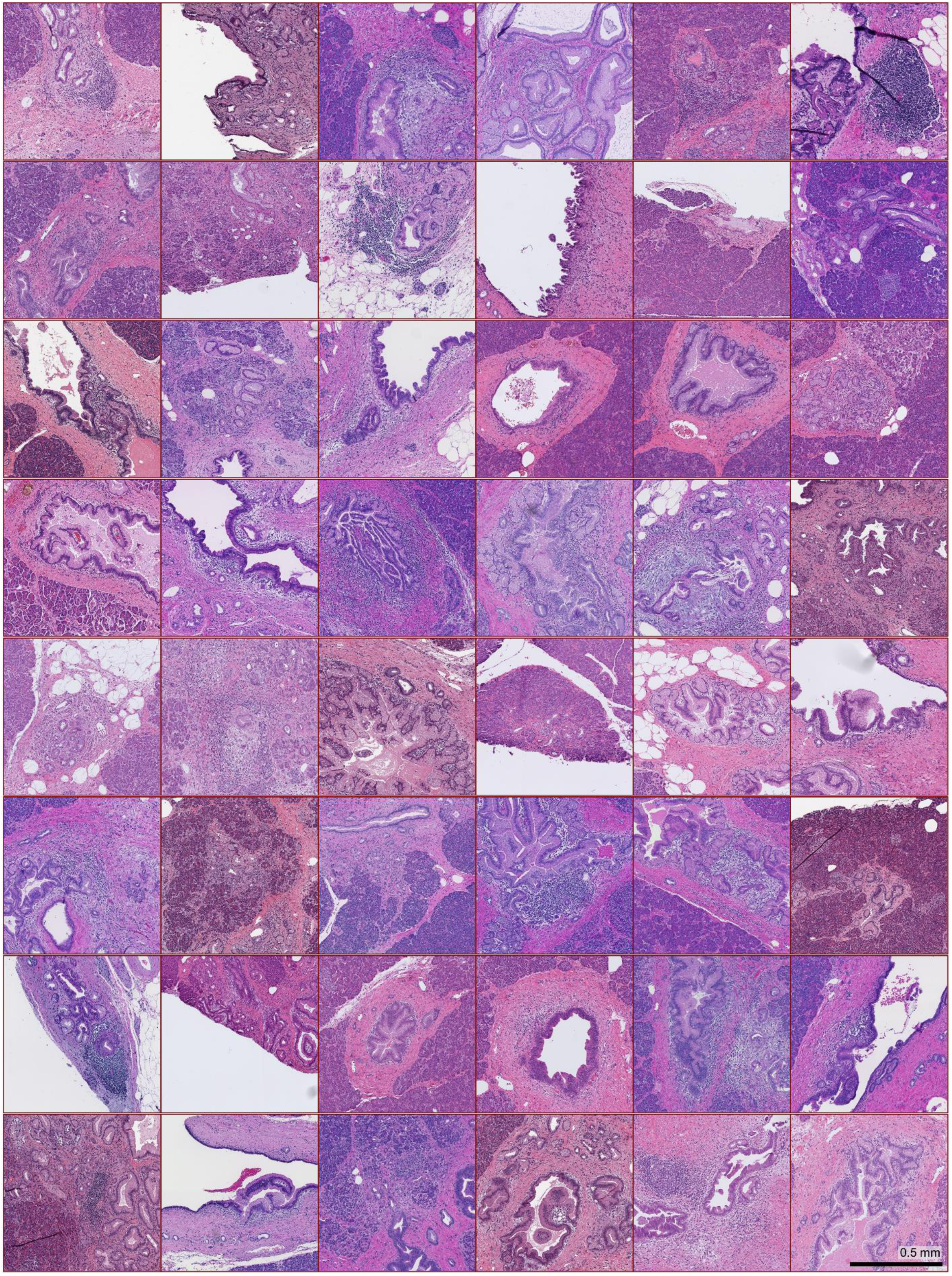
Immune hotspot histology. Sample histology from each of the 48 3D tissue samples containing the immunologically hottest PanIN region of interest in the analyzed sample.

**Fig S5.**
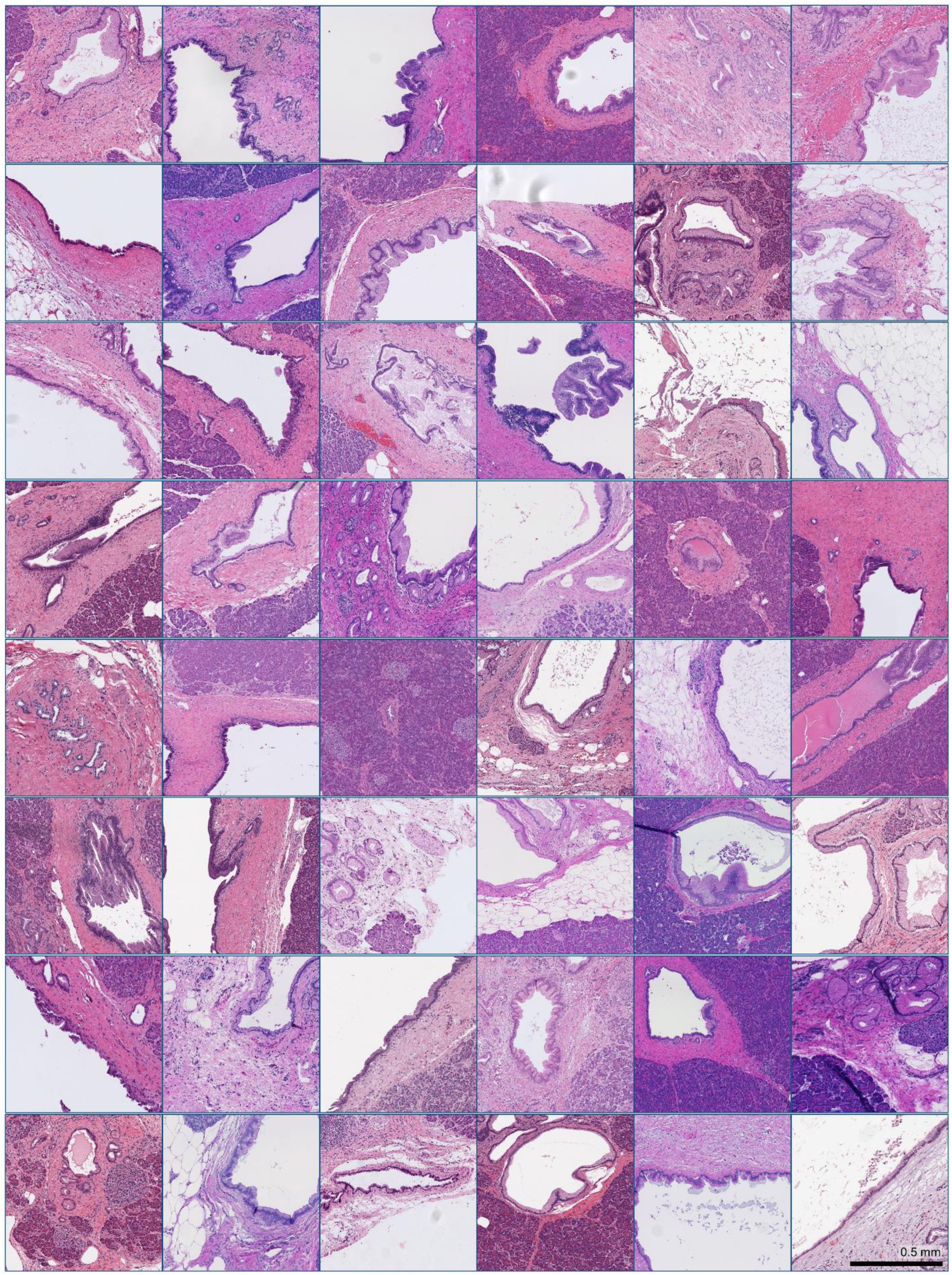
Immune cold spot histology. Sample histology from each of the 48 3D tissue samples containing the immunologically coldest PanIN region of interest in the analyzed sample.

**Fig S6.**
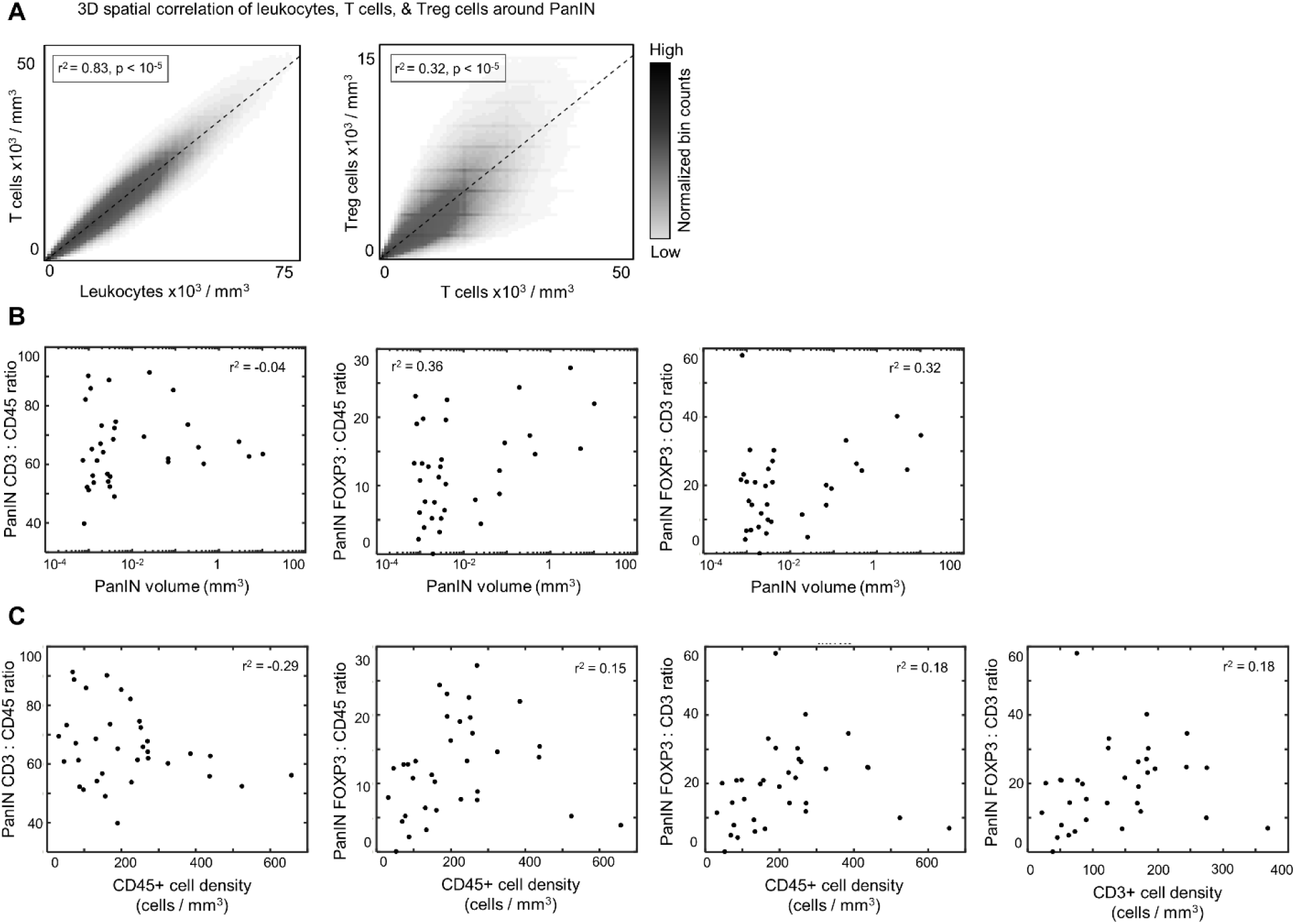
Detailed graphs comparing 3D immune cell compositions around PanIN. (**a**) Pixel-wise correlation between local CD45+ cell density and CD3+cell density, and between CD3+ cell density and FOXP3+ cell density reveals strong correlations (r^2^ = 0.83 and r^2^ = 0.32, respectively). (**b**) Scatter plots depicting the relationship between PanIN volume and immune cell ratios between CD3 and CD45, FOXP3 and CD45, and FOXP3 and CD3. (**c**) Scatter plots depicting the relationship between: CD45+ cell density and immune cell ratios of CD3 to CD45, FOXP3 to CD45, and FOXP3 to CD3, and CD3+ cell density and the immune cell ratio of FOXP3 to CD3.

**Table S1.** Details of imaging mass cytometry antibody cocktail.

| Channel | Antigen | Clone | Dilution | Vendor | Custom |
| --- | --- | --- | --- | --- | --- |
| 89 | CD45 | D9M81 | 125 | CST | X |
| 96-104 | Ruthenium-Tissue Counterstain |  |  |  |  |
| 113 | Collagen | E8F4L | 250 | CST | X |
| 115 | E-cadherin | 24E10 | 125 | CST | X |
| 141 | SMA | 1A4 | 500 | Standard BioTools |  |
| 142 | Podoplanin | D2-40 | 125 | Biolegend | X |
| 143 | VIM | D21H3 | 500 | Standard BioTools |  |
| 144 | CD14 | EPR3653 | 125 | Standard BioTools |  |
| 145 | CD45RO | UCHL1 | 250 | Biolegend | X |
| 146 | CD16 | EPR16784 | 100 | Standard BioTools |  |
| 147 | CD163 | EDHu-1 | 125 | Standard BioTools |  |
| 148 | CK | C11 | 125 | Standard BioTools |  |
| 149 | CD137 | D2Z4Y | 250 | CST | X |
| 150 | PDL1 | E1L3N | 125 | CST | X |
| 151 | PD1 | D4W2J | 125 | BST | X |
| 152 | CD57 | NK/804 | 250 | Standard BioTools |  |
| 153 | Tox/Tox2 | E6I3Q | 250 | CST | X |
| 154 | DC-LAMP | 1010E1.01 (Novus) | 125 | Novus | X |
| 155 | FOXP3 | PCH101 | 75 | Standard BioTools |  |
| 156 | CD4 | EPR6855 | 125 | Standard BioTools |  |
| 158 | pSTAT3 | D3A7 (CST) | 62.5 | CST | X |
| 159 | CD68 | KP1 | 75 | Standard BioTools |  |
| 160 | CD138 | IHC138 (CST) | 125 | CST | X |
| 161 | CD20 | H1 | 125 | Standard BioTools |  |
| 162 | CD8 | C8/144B | 250 | Standard BioTools |  |
| 163 | CD21 | Bu32 (Biolegend) | 250 | Biolegend | X |
| 164 | ARG1 | D4E3M | 75 | Standard BioTools |  |
| 165 | CD33 | 44M12D3 (LS Bio) | 125 | LS Bio | X |
| 166 | CD45RA | HI100 | 250 | Standard BioTools |  |
| 167 | GZMB | D6E9W (CST) | 125 | CST | X |
| 168 | KI67 | B56 | 250 | Standard BioTools |  |
| 170 | CD3 | Polyclonal, C-terminal | 125 | Standard BioTools |  |
| 171 | LAG3 | 17B4 (Novus) | 125 | Novus | X |
| 172 | CD15 | W6D3 | 100 | Biolegend | X |
| 173 | DC-SIGN | DCN46 (IonPath) | 62.5 | Standard BioTools |  |
| 174 | HLADR | LN3 | 250 | Standard BioTools |  |
| 175 | CD86 | E2G8P (CST) | 125 | CST | X |
| 176 | CD206 | E2L9N (CST) | 125 | CST | X |
| 191 | Iridium-DNA |  |  |  |  |
| 193 | Iridium-DNA |  |  |  |  |
| 195 | IMC Seg | 1A36 | 250 | Standard BioTools |  |
| 196 | IMC Seg | 1A37 | 250 | Standard BioTools |  |
| 198 | IMC Seg | 1A38 | 250 | Standard BioTools |  |

## REFERENCES

1. Siegel, R.L., Miller, K.D., Fuchs, H.E., and Jemal, A. (2021). Cancer Statistics, 2021. CA Cancer J Clin 71, 7–33. 10.3322/caac.21654.

2. Morton, D.L., Eilber, F.R., Malmgren, R.A., and Wood, W.C. (1970). Immunological Factors Which Influence Response to Immunotherapy in Malignant Melanoma. Surgery 68, 158-&.

3. Doroshow, D.B., Sanmamed, M.F., Hastings, K., Politi, K., Rimm, D.L., Chen, L.P., Melero, I., Schalper, K.A., and Herbst, R.S. (2019). Immunotherapy in Non-Small Cell Lung Cancer: Facts and Hopes. Clinical Cancer Research 25, 4592–4602. 10.1158/1078-0432.Ccr-18-1538.

4. Herbst, R.S., Morgensztern, D., and Boshoff, C. (2018). The biology and management of non-small cell lung cancer. Nature 553, 446–454. 10.1038/nature25183.

5. Chen, R., Zinzani, P.L., Fanale, M.A., Armand, P., Johnson, N.A., Brice, P., Radford, J., Ribrag, V., Molin, D., Vassilakopoulos, T.P., et al. (2017). Phase II Study of the Efficacy and Safety of Pembrolizumab for Relapsed/Refractory Classic Hodgkin Lymphoma. Journal of Clinical Oncology 35, 2125-+. 10.1200/Jco.2016.72.1316.

6. Bear, A.S., Vonderheide, R.H., and O’Hara, M.H. (2020). Challenges and Opportunities for Pancreatic Cancer Immunotherapy. Cancer Cell 38, 788–802. 10.1016/j.ccell.2020.08.004.

7. Foley, K., Kim, V., Jaffee, E., and Zheng, L. (2016). Current progress in immunotherapy for pancreatic cancer. Cancer Lett 381, 244–251. 10.1016/j.canlet.2015.12.020.

8. Johnston, A.C., Alicea, G.M., Lee, C.C., Patel, P.V., Hanna, E.A., Vaz, E., Forjaz, A., Wan, Z., Nair, P.R., Lim, Y., et al. (2024). Engineering self-propelled tumor-infiltrating CAR T cells using synthetic velocity receptors. bioRxiv. 10.1101/2023.12.13.571595.

9. Yoon, J.H., Jung, Y.J., and Moon, S.H. (2021). Immunotherapy for pancreatic cancer. World J Clin Cases 9, 2969–2982. 10.12998/wjcc.v9.i13.2969.

10. Yadav, D., and Lowenfels, A.B. (2013). The epidemiology of pancreatitis and pancreatic cancer. Gastroenterology 144, 1252–1261. 10.1053/j.gastro.2013.01.068.

11. Bell; A.; Mitchell; J.; Kiemen, A.L., Fujikura, K., Fedor, H., Gambichler, B., Deshpande, A., Wu, P.-H., Sidiropoulos, D.N., Erbe, R., Stern, J., Chan, R., et al. (2022). PanIN and CAF Transitions in Pancreatic Carcinogenesis Revealed with Spatial Data Integration. bioRxiv, in press, Cell Systems. 10.1101/2022.07.16.500312.

12. Zheng, L., Xue, J., Jaffee, E.M., and Habtezion, A. (2013). Role of immune cells and immune-based therapies in pancreatitis and pancreatic ductal adenocarcinoma. Gastroenterology 144, 1230–1240. 10.1053/j.gastro.2012.12.042.

13. Steele, N.G., Biffi, G., Kemp, S.B., Zhang, Y., Drouillard, D., Syu, L., Hao, Y., Oni, T.E., Brosnan, E., Elyada, E., et al. (2021). Inhibition of Hedgehog Signaling Alters Fibroblast Composition in Pancreatic Cancer. Clin Cancer Res 27, 2023–2037. 10.1158/1078-0432.CCR-20-3715.

14. Steele, N.G., Carpenter, E.S., Kemp, S.B., Sirihorachai, V.R., The, S., Delrosario, L., Lazarus, J., Amir, E.D., Gunchick, V., Espinoza, C., et al. (2020). Multimodal Mapping of the Tumor and Peripheral Blood Immune Landscape in Human Pancreatic Cancer. Nat Cancer 1, 1097–1112. 10.1038/s43018-020-00121-4.

15. Liu, C.Y., Xu, J.Y., Shi, X.Y., Huang, W., Ruan, T.Y., Xie, P., and Ding, J.L. (2013). M2-polarized tumor-associated macrophages promoted epithelial-mesenchymal transition in pancreatic cancer cells, partially through TLR4/IL-10 signaling pathway. Lab Invest 93, 844–854. 10.1038/labinvest.2013.69.

16. Zambirinis, C.P., Pushalkar, S., Saxena, D., and Miller, G. (2014). Pancreatic cancer, inflammation, and microbiome. Cancer J 20, 195–202. 10.1097/PPO.0000000000000045.

17. Hiraoka, N., Onozato, K., Kosuge, T., and Hirohashi, S. (2006). Prevalence of FOXP3+ regulatory T cells increases during the progression of pancreatic ductal adenocarcinoma and its premalignant lesions. Clin Cancer Res 12, 5423–5434. 10.1158/1078-0432.CCR-06-0369.

18. Braxton, A.M., Kiemen, A.L., Grahn, M.P., Forjaz, A., Parksong, J., Mahesh Babu, J., Lai, J., Zheng, L., Niknafs, N., Jiang, L., et al. (2024). 3D genomic mapping reveals multifocality of human pancreatic precancers. Nature. 10.1038/s41586-024-07359-3.

19. Matsuda, Y., Furukawa, T., Yachida, S., Nishimura, M., Seki, A., Nonaka, K., Aida, J., Takubo, K., Ishiwata, T., Kimura, W., et al. (2017). The Prevalence and Clinicopathological Characteristics of High-Grade Pancreatic Intraepithelial Neoplasia: Autopsy Study Evaluating the Entire Pancreatic Parenchyma. Pancreas 46, 658–664. 10.1097/MPA.0000000000000786.

20. Carpenter, E.S., Elhossiny, A.M., Kadiyala, P., Li, J., McGue, J., Griffith, B.D., Zhang, Y., Edwards, J., Nelson, S., Lima, F., et al. (2023). Analysis of Donor Pancreata Defines the Transcriptomic Signature and Microenvironment of Early Neoplastic Lesions. Cancer Discov 13, 1324–1345. 10.1158/2159-8290.CD-23-0013.

21. Siegel, R.L., Miller, K.D., Wagle, N.S., and Jemal, A. (2023). Cancer statistics, 2023. CA Cancer J Clin 73, 17–48. 10.3322/caac.21763.

22. Lapidaire, W., Forkert, N.D., Williamson, W., Huckstep, O., Tan, C.M., Alsharqi, M., Mohamed, A., Kitt, J., Burchert, H., Mouches, P., et al. (2023). Aerobic exercise increases brain vessel lumen size and blood flow in young adults with elevated blood pressure. Secondary analysis of the TEPHRA randomized clinical trial. Neuroimage Clin 37, 103337. 10.1016/j.nicl.2023.103337.

23. Society, A.C. Cancer Facts & Figures 2022.

24. Kiemen, A.L., Braxton, A.M., Grahn, M.P., Han, K.S., Babu, J.M., Reichel, R., Jiang, A.C., Kim, B., Hsu, J., Amoa, F., et al. (2022). CODA: quantitative 3D reconstruction of large tissues at cellular resolution. Nature Methods 19, 1490–1499. 10.1038/s41592-022-01650-9.

25. Kiemen, A.L.D., L.; Shen, Y; Zhu, Y.; Matos-Romero, V.; Forjaz, A.; Campbell, K.; Dhana, W.; Cornish, T.; Braxton, A.; Wu, P.; Fishman, E.; Wood, L.; Wirtz, D.; Hruban, R. (2024). PanIN or IPMN? Redefining lesion size in three dimensions. American Journal of Surgical Pathology.

26. Kiemen, A.L., Wu, P.H., Braxton, A.M., Cornish, T.C., Hruban, R.H., Wood, L., Wirtz, D., and Zwicker, D. (2024). Power-law growth models explain incidences and sizes of pancreatic cancer precursor lesions and confirm spatial genomic findings. Science Advances. 10.1101/2023.12.01.569633.

27. Forjaz, A.V., E. Matos-Romero, V.; Joshi, S.; Fujikara, K.; Braxton, A.M.; Jiang, A.; Cornish, T.; Hong, S.M.; Hruban, R.H.; Wood, L.; Wu, P.H.; Kiemen, A.; Wirtz, D. (2023). Three-dimensional assessments are necessary to determine the true spatial tissue composition of diseased tissues. Biorxiv.

28. Kiemen, A.L., Forjaz, A., Sousa, R., Han, K.S., Hruban, R.H., Wood, L.D., Wu, P.H., and Wirtz, D. (2024). High-Resolution 3D Printing of Pancreatic Ductal Microanatomy Enabled by Serial Histology. Adv Mater Technol-Us 9. 10.1002/admt.202301837.

29. Deshpande, A.L., M.; Sidiripoulos, D. N.; Zhangm S.; Yuanm L; Bell, A.; Zhu, Q. Jin Ho, W.; Santa-Maria, C.; Gilkes, D.; Williams, S. R.; Uytingco, C.R.; Chew, J.; Hartnett, A.; Bent, Z.W.; Favorov, A. V.; Popel, A.S.; Yarchoan, M.; Kiemen, A.; Wu, P.H.; Fujikura, K.; Wirtz, D.; Wood, L.; Zheng, L.; Jaffee, E. M.; Anders, R.; Danilova, L.; Stein-O’Brien, G.; Kagohara, L.T.; Fertig, E. (2023). Uncovering the spatial landscape of molecular interactions within the tumor microenvironment through latent spaces. Cell Systems.

30. Mi, H., Gong, C., Sulam, J., Fertig, E.J., Szalay, A.S., Jaffee, E.M., Stearns, V., Emens, L.A., Cimino-Mathews, A.M., and Popel, A.S. (2020). Digital Pathology Analysis Quantifies Spatial Heterogeneity of CD3, CD4, CD8, CD20, and FoxP3 Immune Markers in Triple-Negative Breast Cancer. Front Physiol 11, 583333. 10.3389/fphys.2020.583333.

31. Lin, J.R., Wang, S., Coy, S., Chen, Y.A., Yapp, C., Tyler, M., Nariya, M.K., Heiser, C.N., Lau, K.S., Santagata, S., and Sorger, P.K. (2023). Multiplexed 3D atlas of state transitions and immune interaction in colorectal cancer. Cell 186, 363–381 e319. 10.1016/j.cell.2022.12.028.

32. Tian, J., Qian, B., Zhang, S., Guo, R., Zhang, H., Jeannon, J.P., Jin, R., Feng, X., Zhan, Y., Liu, J., et al. (2020). Three-dimensional reconstruction of laryngeal cancer with whole organ serial immunohistochemical sections. Sci Rep 10, 18962. 10.1038/s41598-020-76081-7.

33. Yagi, Y., Aly, R.G., Tabata, K., Barlas, A., Rekhtman, N., Eguchi, T., Montecalvo, J., Hameed, M., Manova-Todorova, K., Adusumilli, P.S., and Travis, W.D. (2020). Three-Dimensional Histologic, Immunohistochemical, and Multiplex Immunofluorescence Analyses of Dynamic Vessel Co-Option of Spread Through Air Spaces in Lung Adenocarcinoma. J Thorac Oncol 15, 589–600. 10.1016/j.jtho.2019.12.112.

34. Arganda-Carreras, I., Fernandez-Gonzalez, R., Munoz-Barrutia, A., and Ortiz-De-Solorzano, C. (2010). 3D reconstruction of histological sections: Application to mammary gland tissue. Microsc Res Tech 73, 1019–1029. 10.1002/jemt.20829.

35. Lotz, J.M., Hoffmann, F., Lotz, J., Heldmann, S., Trede, D., Oetjen, J., Becker, M., Ernst, G., Maas, P., Alexandrov, T., et al. (2017). Integration of 3D multimodal imaging data of a head and neck cancer and advanced feature recognition. Biochim Biophys Acta Proteins Proteom 1865, 946–956. 10.1016/j.bbapap.2016.08.018.

36. Groot, A.E.d., Myers, K.V., Krueger, T.E.G., Kiemen, A.L., Nagy, N.H., Brame, A., Torres, V.E., Zhang, Z., Trabzonlu, L., Brennen, W.N., et al. (2021). Characterization of tumor-associated macrophages in prostate cancer transgenic mouse models. The Prostate. John Wiley & Sons, Ltd.

37. Kiemen, A.L., Choi, Y., Braxton, A.M., Almagro Perez, C., Graham, S., Grahn, M.P., Nanda, N., Roberts, N., Wood, L., Wu, P., et al. (2022). Intraparenchymal metastases as a cause for local recurrence of pancreatic cancer. Histopathology. 10.1111/his.14839.

38. Crawford, A.J., Forjaz, A., Bhorkar, I., Roy, T., Schell, D., Queiroga, V., Ren, K., Kramer, D., Bons, J., Huang, W., et al. (2023). Precision-engineered biomimetics: the human fallopian tube. bioRxiv. 10.1101/2023.06.06.543923.

39. Lee, M.H., Russo, G.C., Rahmanto, Y.S., Du, W.X., Crawford, A.J., Wu, P.H., Gilkes, D., Kiemen, A., Miyamoto, T., Yu, Y., et al. (2022). Multi-compartment tumor organoids. Mater Today 61, 104–116. 10.1016/j.mattod.2022.07.006.

40. Chen, L.C., Papandreou, G., Kokkinos, I., Murphy, K., and Yuille, A.L. (2018). DeepLab: Semantic Image Segmentation with Deep Convolutional Nets, Atrous Convolution, and Fully Connected CRFs. IEEE Transactions on Pattern Analysis and Machine Intelligence. IEEE Computer Society.

41. Daniluk, J., Liu, Y., Deng, D., Chu, J., Huang, H., Gaiser, S., Cruz-Monserrate, Z., Wang, H., Ji, B., and Logsdon, C.D. (2012). An NF-kappaB pathway-mediated positive feedback loop amplifies Ras activity to pathological levels in mice. J Clin Invest 122, 1519–1528. 10.1172/JCI59743.

42. Collins, M.A., Yan, W., Sebolt-Leopold, J.S., and di Magliano, M.P. (2014). MAPK Signaling Is Required for Dedifferentiation of Acinar Cells and Development of Pancreatic Intraepithelial Neoplasia in Mice. Gastroenterology 146, 822-+. 10.1053/j.gastro.2013.11.052.

43. Farrow, B., Sugiyama, Y., Chen, A., Uffort, E., Nealon, W., and Evers, M. (2004). Inflammatory mechanisms contributing to pancreatic cancer development. Annals of Surgery 239, 763–769. 10.1097/01.sla.0000128681.76786.07.

44. Medzhitov, R. (2008). Origin and physiological roles of inflammation. Nature 454, 428–435. 10.1038/nature07201.

45. LeBlanc, J.K., Chen, J.H., Al-Haddad, M., Luz, L., McHenry, L., Sherman, S., Juan, M., and Dewitt, J. (2014). Can Endoscopic Ultrasound Predict Pancreatic Intraepithelial Neoplasia Lesions in Chronic Pancreatitis? on. Pancreas 43, 849–854. Doi 10.1097/Mpa.0000000000000142.

46. Tirkes, T., Yadav, D., Conwell, D.L., Territo, P.R., Zhao, X., Persohn, S.A., Dasyam, A.K., Shah, Z.K., Venkatesh, S.K., Takahashi, N., et al. (2022). Quantitative MRI of chronic pancreatitis: results from a multi-institutional prospective study, magnetic resonance imaging as a non-invasive method for assessment of pancreatic fibrosis (MINIMAP). Abdom Radiol 47, 3792–3805. 10.1007/s00261-022-03654-7.

47. Bieliuniene, E., Frokjær, J.B., Pockevicius, A., Kemesiene, J., Lukosevicius, S., Basevicius, A., Barauskas, G., Dambrauskas, Z., and Gulbinas, A. (2019). Magnetic Resonance Imaging as a Valid Noninvasive Tool for the Assessment of Pancreatic Fibrosis. Pancreas 48, 85–93. 10.1097/Mpa.0000000000001206.

48. Klöppel, G., Detlefsen, S., and Feyerabend, B. (2004). Fibrosis of the pancreas:: the initial tissue damage and the resulting pattern. Virchows Arch 445, 1–8. 10.1007/s00428-004-1021-5.

49. Zaheer, A., Singh, V.K., Akshintala, V.S., Kawamoto, S., Tsai, S.D., Gage, K.L., and Fishman, E.K. (2014). Differentiating Autoimmune Pancreatitis From Pancreatic Adenocarcinoma Using Dual-Phase Computed Tomography. J Comput Assist Tomo 38, 146–152. 10.1097/RCT.0b013e3182a9a431.

50. Brune, K., Abe, T., Canto, M., O’Malley, L., Klein, A.P., Maitra, A., Adsay, N.V., Fishman, E.K., Cameron, J.L., Yeo, C.J., et al. (2006). Multifocal neoplastic precursor lesions associated with lobular atrophy of the pancreas in patients having a strong family history of pancreatic cancer. Am J Surg Pathol 30, 1067–1076.

51. Ashley L. Kiemen, L.D., Yu Shen, Yutong Zhu, Valentina Matos-Romero, André Forjaz, Kurtis Campbell, Will Dhana, Toby Cornish, Alicia M. Braxton, PeiHsun Wu, Elliot K. Fishman, Laura D. Wood, Denis Wirtz, Ralph Hruban (2024). PanIN or IPMN? Redefining lesion size in three dimensions. Am J Surg Pathol.

52. Kiemen, A.L., Dbouk, M., Diwan, E.A., Forjaz, A., Dequiedt, L., Baghdadi, A., Madani, S.P., Grahn, M.P., Jones, C., Vedula, S., et al. (2024). Magnetic Resonance Imaging-Based Assessment of Pancreatic Fat Strongly Correlates With Histology-Based Assessment of Pancreas Composition. Pancreas 53, e180–e186. 10.1097/MPA.0000000000002288.

53. Goode, A., Gilbert, B., Harkes, J., Jukic, D., and Satyanarayanan, M. (2013). OpenSlide: A vendor-neutral software foundation for digital pathology. Journal of Pathology Informatics. Wolters Kluwer --Medknow Publications.

54. Park, H.S., and Jun, C.H. (2009). A simple and fast algorithm for K-medoids clustering. Expert Syst Appl 36, 3336–3341. 10.1016/j.eswa.2008.01.039.

